# Chemical propulsion of hemozoin crystal motion in malaria parasites

**DOI:** 10.1101/2025.04.25.650681

**Authors:** Erica M. Hastings, Tomasz Skora, Keith R. Carney, Henry C. Fu, Tamara C. Bidone, Paul A. Sigala

## Abstract

Malaria parasites infect red blood cells where they digest host hemoglobin and release free heme inside a lysosome-like organelle called the food vacuole. To detoxify excess heme, parasites form hemozoin crystals that rapidly tumble inside this compartment. Hemozoin formation is critical for parasite survival and central to antimalarial drug activity. Although the static structural properties of hemozoin have been extensively investigated, crystal motion and its underlying mechanism have remained puzzling. We used quantitative image analysis to determine the timescale of motion, which requires the intact vacuole but does not require the parasite itself. Using single particle tracking and Brownian dynamics simulations with experimentally derived interaction potentials, we found that hemozoin motion exhibits unexpectedly tight confinement but is much faster than thermal diffusion. Hydrogen peroxide, which is generated at high levels in the food vacuole, has been shown to stimulate the motion of synthetic metallic nanoparticles via surface-catalyzed peroxide decomposition that generates propulsive kinetic energy. We observed that peroxide stimulated the motion of isolated crystals in solution and that conditions that suppress peroxide formation slowed hemozoin motion inside parasites. These data suggest that surface-exposed metals on hemozoin catalyze peroxide decomposition to drive crystal motion. This work reveals hemozoin motion in malaria parasites as a biological example of an endogenous self-propelled nanoparticle. This mechanism of propulsion likely serves a physiological role to reduce oxidative stress to parasites from hydrogen peroxide produced by large-scale hemoglobin digestion during blood-stage infection.

**Significance statement:** Hemozoin crystal formation is a major antimalarial drug target as it is essential for survival of *Plasmodium* parasites that cause malaria, one of the world’s most devastating infectious diseases. Hemozoin crystals rapidly tumble inside the parasite food vacuole, and the mechanism and significance of this motion have been mysterious. Using quantitative live-cell imaging and computational modeling, we discovered that hemozoin motion is driven by catalytic decomposition of hydrogen peroxide on the crystal surface, a mechanism analogous to synthetic nanomotors. Hemozoin crystals have been viewed as inert detoxification products. Our work reframes the physiological role of these crystals as catalytically active nanoparticles whose mechanism of propulsion helps to neutralize toxic hydrogen peroxide generated by parasite digestion of hemoglobin during blood-stage infection.

## Introduction

Malaria remains a deadly infectious disease that kills over 600,000 people each year and is caused by *Plasmodium* parasites that invade and destroy red blood cells (RBCs) (1). During blood-stage infection, parasites digest most of the RBC hemoglobin within an acidic food vacuole (FV) that is similar to a mammalian lysosome (2). Hemoglobin proteolysis releases large quantities of the iron-containing cofactor, heme, which parasites detoxify by forming multiple ∼500 nm brick-shaped hemozoin crystals within the ∼1 μm-radius FV (**Fig. 1A**) (3, 4). The biocrystallization of heme is a critical parasite survival strategy and major antimalarial drug target, such that free heme that escapes incorporation into crystals is thought to underpin chloroquine activity and serve as the key activator of frontline artemisinin therapies (5, 6).

**Figure 1:**
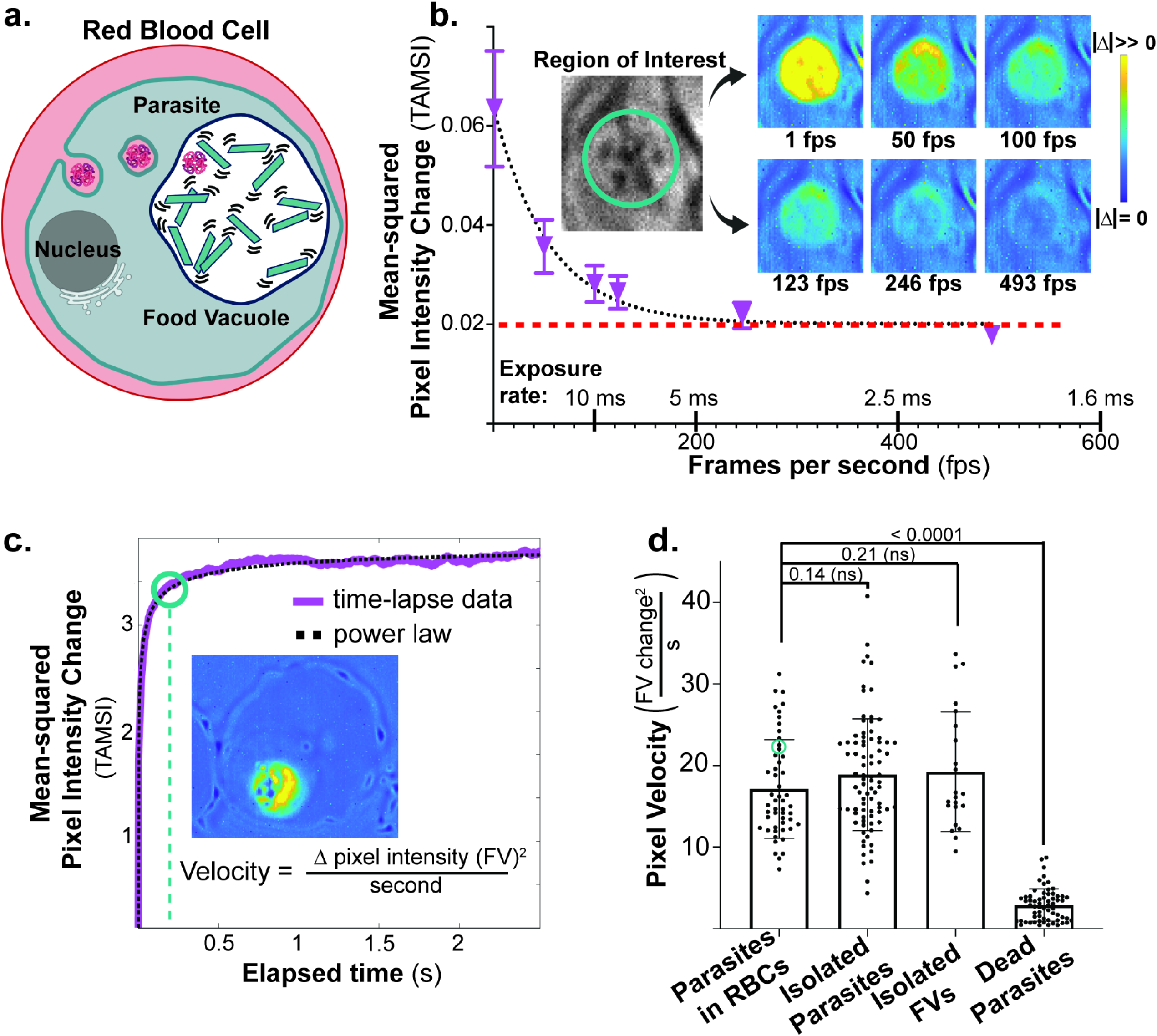
Quantification of hemozoin motion and its dependence on the FV environment. **a**, Diagram of a parasite-infected RBC, trafficking of host hemoglobin to the food vacuole, and formation of moving hemozoin crystals. **b**, Parasite FV regions of interest (inset) were imaged for variable exposure times. The pixel intensity change (Δ) in the region of interest was plotted versus time differences between frames. Heat map images (inset) show the gradient of pixel intensity changes, with yellow pixels indicating largest change and dark blue pixels indicating smallest change. Pixel intensity change dependence on frame rate was empirically fit to an exponential decay equation (black dotted line). Red dashed line indicates lower limit of background camera detection. **c**, Plots of time-averaged mean squared pixel intensity change (TAMSI) within the FV region as a function of elapsed time revealed a plateau (decorrelation) beyond 0.2 s. Data were fit with a power-law growth function, provided in Methods, to determine the average time to plateau. An apparent pixel velocity was determined by dividing the TAMSI value at the average plateau time of 0.18 s in native parasites (teal circle). **d**, Population analysis of pixel velocities for FVs in live, fractionated, or dead (drug-treated) parasites. Teal circle shows pixel velocity of FV analyzed in Fig. 1c. Dead parasites were treated with a lethal dose of chloroquine (shown) or other antimalarial drugs. Hemozoin motion stopped in all treatments. Statistical analysis was done by Student’s t-test to determine the indicated p-values. Error bars represent the standard deviation of the mean.

Within the FV organelle, hemozoin crystals dynamically tumble in a rapid and unpredictable fashion (***SI Appendix,* Movie S1**) (7-10). Although ubiquitous, these dynamics are sparsely studied and have been described as Brownian in nature (7, 8). However, prior chemical and genetic perturbations of the FV have provided evidence that hemozoin motion can be stopped, suggesting additional energy inputs beyond thermal diffusion (***SI Appendix,* Movie S2, Fig. S1A**) (9-11). Previous studies also provide conflicting evidence about whether hemozoin tumbling requires a physiologically active cell and thus serves as a biomarker of parasite viability or responds to local properties of the FV environment independent of the parasite (7-10). Therefore, uncovering the physical nature and causative mechanism of crystal dynamics is key to unlocking a full understanding of hemozoin and its critical function in malaria parasite survival and drug susceptibility.

## Results

### Hemozoin tumbling occurs on ms timescale

The rapid movement of hemozoin crystals is a hallmark of RBC infection by *P. falciparum*, which is the most virulent parasite species in Africa and causes nearly all human malaria deaths. To determine the timescale of hemozoin dynamics within the FV, we imaged *P. falciparum*-infected RBCs with brightfield microscopy using progressively faster frame rates (***SI Appendix,* Fig. S1B**). We observed that the pixel intensity change between consecutive frames within the FV region decreased with increasing frame rate until plateauing at ∼200 frames per second (Δt = 5 ms) (**Fig. 1B**). This observation indicates that the fastest detectable displacements of hemozoin crystals occur on a 5 ms timescale. Therefore, we used a frame rate of ≥200 frames per second for all subsequent imaging.

### Hemozoin motion is a local FV property

To better understand the nature and origin of hemozoin motion within the FV, we sought to quantify the movement of individual crystals. Single-particle tracking (SPT) was unsuccessful due to crystal congestion and spatially overlapping trajectories in live parasites. To overcome this challenge, we developed an alternative method that quantified the mean squared pixel intensity change (TAMSI) within the FV as a function of elapsed time, which we define as “pixel velocity” (**Fig. 1C, *SI Appendix,* Movie S3, Fig. S2A-C**). This analysis revealed >10-fold differences between hemozoin motion in live versus dead (drug-inactivated) parasites, suggesting that the physiological condition of the FV underpins crystal dynamics (**Fig. 1D, *SI Appendix,* Fig. S2D**). To further probe the biological requirements for hemozoin motion, we fractionated parasites and tested various conditions for crystal motion. We measured hemozoin crystal TAMSI in fractionated intact parasites and isolated FVs and observed nearly identical pixel velocities across all conditions except dead parasites (**Fig. 1D, *SI Appendix,* Fig. S2A-D**). These findings indicate that hemozoin motion is a local property of the biochemically active FV environment and independent of the parasite’s cellular context. However, this motion stops in lethally drug-treated parasites, suggesting a loss in FV membrane integrity and/or physiological activity in dead parasites.

### Single-particle tracking of crystal motion

To overcome FV congestion and enable accurate single-particle tracking (SPT) analysis of crystal displacements, we expanded the vacuoles using hypotonic conditions previously observed to enlarge mammalian lysosomes (12-14). This expansion increased FV radii from ∼1 μm to ∼3 μm while preserving the rotational and translational components of hemozoin crystal motion (**Fig. 2A, *SI Appendix,* Fig. S3A, Movie S4**). Critically, the trajectories of individual crystals within expanded FVs could be reliably tracked for up to half a second (≥ 100 frames) to enable detailed analyses of their motion (**Fig. 2A**).

**Figure 2:**
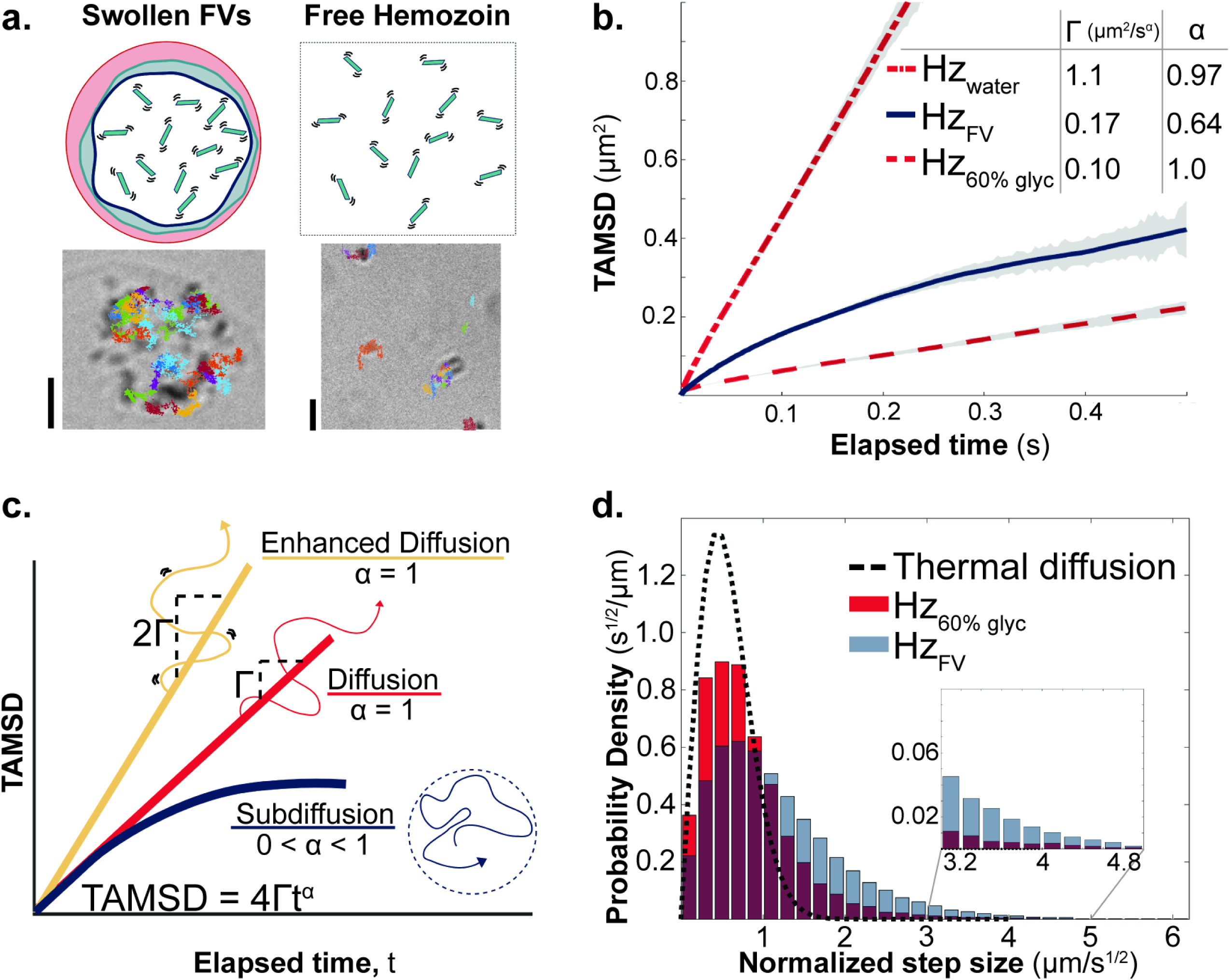
Expanded food vacuoles enable single-particle tracking and reveal enhanced but confined motion. **a**, Hypotonic conditions increase FV radius by ≥2-fold while maintaining crystal motion but avoiding crystal overlap (**Movie S4**). FV crystals are compared to isolated crystals in media with defined viscosities. Individual particle tracking showing longest tracks. Scale bars = 2 μm. **b**, Time-averaged mean squared displacement (TAMSD) of hemozoin crystals in expanded FVs compared to free hemozoin in matched viscosity and water with fit values from Table S1. Error bars represent the standard deviation of the means of individual trajectories and are shown in grey. **c**, Diffusion profile scheme showing generalized interpretations of TAMSD analysis. Γ is generalized diffusion coefficient (μm^2^/s^α^), α is power law exponent. **d**, Histogram showing probability density of average step size (distance traveled) of each crystal over normalized time. Steps were normalized by dividing by square root time between beginning and end of step (due to varying elapsed times in data set). Maroon bins show overlapping data in probability distributions.

Within expanded FVs, we collected over 3500 trajectories of individual hemozoin crystals and analyzed their time-averaged mean squared displacements (TAMSDs) (***SI Appendix,* Fig. S3C**). The weighted average of these TAMSDs as a function of elapsed time showed an anomalous nonlinear time dependence with a diminishing slope (**Fig. 2B-C**). This observation contradicts predictions for simple diffusion based on the Einstein-Smoluchowski equation, where the TAMSD increases linearly with time (MSD = 4Dt) (15-18). Instead, the TAMSD data for hemozoin crystals in the FV were best fit by a power law (MSD = 4Γt^α^), with an α value <1 indicative of subdiffusion (**Fig. 2C**) (19).

To compare the anomalous motion of FV hemozoin crystals to conditions where purely diffusive motion is expected, we isolated hemozoin crystals and suspended them in aqueous solutions of variable viscosity (**Fig. 2A**). Viscosity within the parasite FV is unknown but expected to be similar to a mammalian lysosome, ∼100 – 400 cP (20-23). Based on these values, we estimated the viscosity of expanded vacuoles due to water influx to be 10 – 40 cP, whose lower limit is equal to the viscosity of 60% w/v glycerol (aq) (***SI Appendix,* Fig. S3B**). As expected, SPT of isolated hemozoin crystals or inert nanobeads in water or 60% glycerol displayed a linear time dependence (α ∼ 1), consistent with pure diffusion, while hemozoin motion within expanded FVs exhibited nonlinear behavior (α = 0.64 ± 0.12) (**Fig. 2B, *SI Appendix,* Table S1, Fig. S3B**).

Based on fits to the power law form of the Einstein-Smoluchowski equation, hemozoin within expanded FVs had a generalized diffusion coefficient (Γ) of ∼ 0.17 μm^2^/s^α^, which was intermediate between that of isolated hemozoin in water (∼ 1.1 μm^2^/s^α^, 1 cP) and in 60% glycerol (∼ 0.10 μm^2^/s^α^, 10 cP). The enhanced (faster) diffusion of hemozoin in FVs compared to 60% glycerol was particularly prominent, given the expectation that both environments have similar viscosities. Possible explanations for this difference in slope include inaccuracy in FV viscosity, which may be less than 10 cP, or the presence of additional factors within the FV that stimulate crystal motion.

To test the accuracy of our estimated viscosity in expanded FVs, we analyzed the distribution of normalized crystal step sizes (μm/s^1/2^) for hemozoin in expanded FVs, 60% glycerol (10 cP), or a thinner solution of 40% glycerol (4 cP). For a spherical particle of radius 0.2 μm, Brownian motion theory predicts a characteristic distribution of particle steps centered around ∼ 0.6 μm/s^1/2^at a viscosity of 10 cP or ∼1.2 µm/s^1/2^ at 4 cP (15-18). We observed a most frequent step size of ∼ 0.6 μm/s^1/2^ for hemozoin in expanded FVs and isolated crystals in 60% glycerol (**Fig. 2D, *SI Appendix,* Fig. S3D-E**). In contrast, isolated crystals in 40% glycerol displayed a recurring step size of ∼1.2 µm/s^1/2^ that was two-fold larger than FV hemozoin (***SI Appendix,* Fig. S3D-E**). The close agreement between peak frequency of hemozoin step sizes within expanded FVs and 60% glycerol (10 cP) and disagreement with the larger average step sizes in 40% glycerol (4 cP) strongly support the conclusion that expanded FVs have an effective viscosity of 10 cP and that the motion of FV hemozoin is faster than predicted for thermal diffusion at this viscosity.

Further dissection of the histograms revealed that hemozoin in expanded FVs exhibited a heavier tail of longer step sizes compared to crystals in 60% glycerol (**Fig. 2D**). To quantify this difference, we attempted to fit both distributions to a single population of moving particles. The 60% glycerol histogram was accurately fit by this model, which indicated a single population of particles with a diffusion coefficient of ∼ 0.17 μm^2^/s. However, the same fit to the histogram of crystals in expanded FVs could not be explained by a single population and instead revealed that the distribution was more accurately decomposed into two populations, with 33% of crystals having an expected diffusion coefficient (D_1_) of 0.13 μm^2^/s and 67% of crystals with a much higher (faster) diffusion coefficient (D_2_) of 0.63 μm^2^/s (***SI Appendix,* Fig. S3E, Table S5**). Since D_1_ is close to the expected D for thermal diffusivity in 60% glycerol, we interpret the first population as purely diffusive particles. The second, faster-moving population for FV hemozoin suggests either that the size variation of hemozoin crystals differed between FVs and the glycerol environment or that an additional energy source transiently accelerates crystal diffusion within the FV. To analyze the potential for size heterogeneity, we solved for the expected particle size at the faster diffusion coefficient. This analysis indicated that most particles would have to be smaller than the diffraction limit of light (radius of ∼ 0.03 μm) to account for faster diffusion and therefore unlikely to be seen or tracked through a brightfield microscope (24). Based on these findings, we conclude that the *Plasmodium* food vacuole viscosity is indeed similar to a mammalian lysosome and that crystal motion deviates from our simplified experimental models of Brownian diffusion with additional features of both confined and stimulated motion.

### Simulated multi-particle crystal motion

Hemozoin crystals within the parasite FV have a cuboidal shape, are confined by the surrounding FV membrane, and interact with each other due to spatial crowding. These biophysical features differ from isolated crystals in solution or simple spheres modeled by the Einstein-Smoluchowski equation for simple Brownian motion. To determine if differences in crystal shape, FV boundary effects, and/or particle-particle interactions contribute to the faster but constrained diffusion of FV hemozoin, we performed stochastic Brownian dynamics simulations (25). These simulations incorporated interaction potentials and boundary conditions designed to mimic expanded FV environments (***SI Appendix,* Fig. S4A**). Specifically, we simulated diffusive motion of both hemozoin-shaped bricks and spherical particles under dimensional confinement conditions similar to those in our experiments and analyzed their trajectories to determine TAMSD values (***SI Appendix,* Movie S5-S6, Table S6**). Cuboid bricks were modeled using surface-covering nodes with volume exclusion enforced by a repulsive Lennard-Jones potential (**Fig. 3A-B**). Simulations of spherical particles included both attractive and repulsive terms of the Lennard-Jones potential. This potential was parameterized from experimental SPT data of FV hemozoin using the Direct Boltzmann Inversion method and exhibited a very shallow minimum of -0.24 kcal/mol (≪ k_B_T) at 0.82 μm, indicating weak attraction between crystals (**Fig. 3B, *SI Appendix,* Fig. S4B-C**) (26).

**Figure 3:**
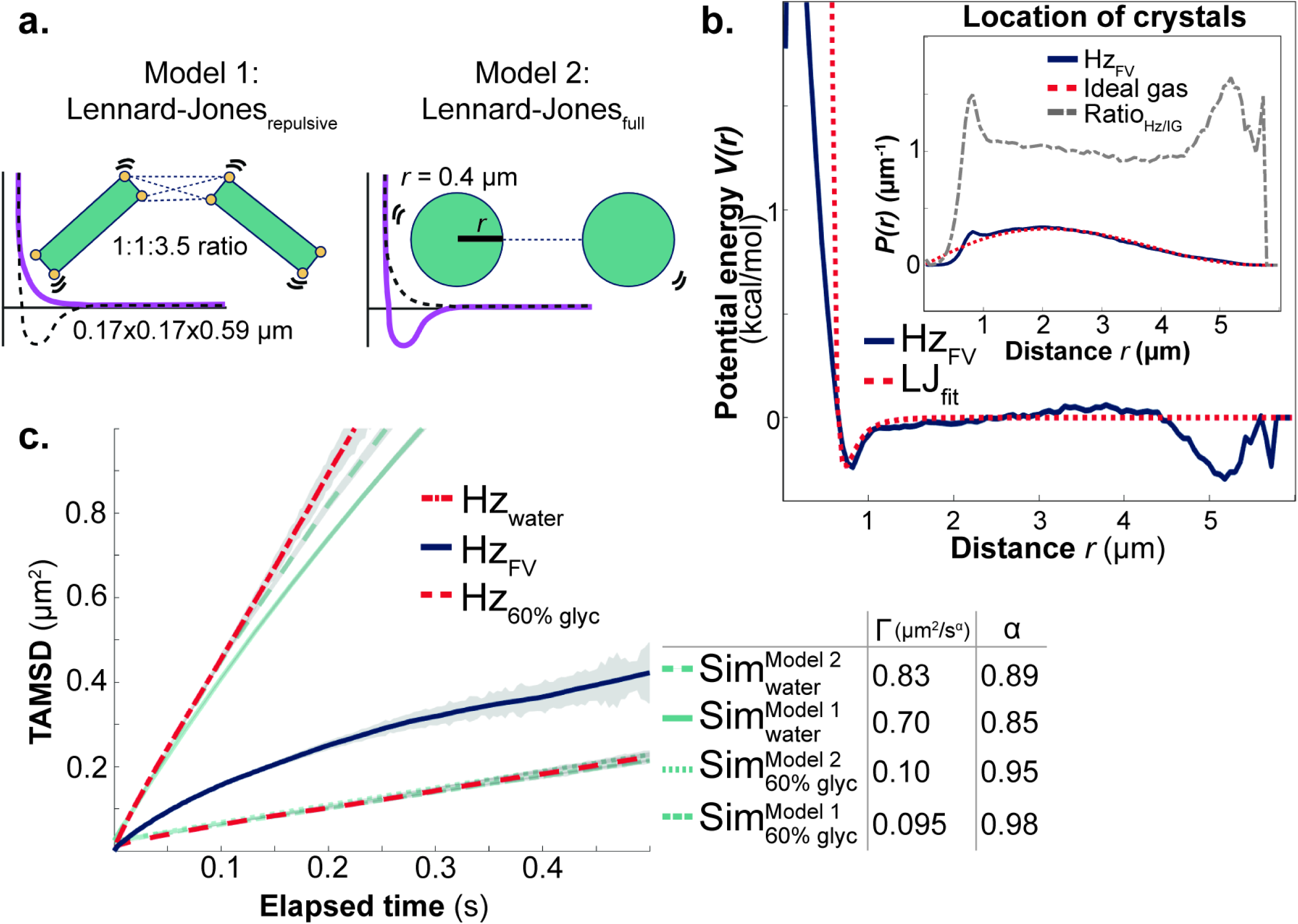
Stochastic Brownian dynamics simulations of hemozoin crystal motion in distinct environments cannot account for anomalous FV features. **a**, Different geometric shapes and particle-particle interactions were used to compare Brownian models. Model 1 used cuboidal bricks with a repulsive Lennard-Jones interaction within a contact distance of 500 nm. Cuboidal shapes used dimensions as shown following previous literature (4, 74). Model 2 used simplified spheres of equal crystal volume with a full Lennard-Jones potential including attraction and repulsion. **b**, Experimental tracks were used to calculate the interaction potentials between crystals. The calculated interaction potential looked like a fully defined Lennard-Jones potential with weak attraction and then repulsion at the minimal distance between two crystals. Model 2 used these interaction potentials. Inset shows probability of crystal-crystal distance in experiment vs. in an ideal gas scenario. **c**, TAMSD of hemozoin crystals in expanded FVs compared to simulations from Models 1 and 2 from Fig. 3A. See Tables S6 - S8 for fitted TAMSD values. Error bars represent the standard deviation of the means of individual trajectories and are shown in grey.

Simulated TAMSDs were adjusted to account for the static localization error in the experimental trajectories resulting from microscope noise that systematically offsets the observed particle position from the true position (27). After making this correction, we observed a strong concordance between the simulated models and experimental SPT analysis of isolated hemozoin in water and 60% glycerol over 0.5 s (**Fig. 3C**). Using the power-law form of the Einstein-Smoluchowski equation to fit the simulated data, we determined α exponents of ≥0.85 and ≥0.98 for crystal motion at viscosities equal to water and 60% glycerol, respectively, despite particle-particle interactions and confinement within a spherical boundary of ∼2.5 μm radius (***SI Appendix,* Tables S6 - S8**). These results closely matched the experimental data for isolated hemozoin in these environments. Simulated TAMSDs in water showed a slight curvature toward subdiffusive behavior, consistent with the expectation that particles reach the simulated boundary sooner in this lower-viscosity environment. Most importantly, the α exponents in simulations were still significantly greater than the α exponent obtained experimentally for FV hemozoin (0.64 ± 0.12) (**Fig. 3C**, ***SI Appendix,* Fig. S4C, Tables S6 – S8**). These findings suggest that differences in hemozoin shape, FV boundary constraints, and interactions between hemozoin crystals do not explain the faster and nonlinear time dependence of FV hemozoin motion.

### Propulsion and micro-confinements explain hemozoin motion

To understand the enhanced diffusivity and constrained TAMSD properties of FV hemozoin compared to isolated crystals, we further analyzed their TAMSD and normalized step-size distribution (**Fig. 2B,D**). The nonlinear shape of the hemozoin TAMSD in FVs aligns with theoretical predictions for particle diffusion where the TAMSD plateaus at ∼ 0.8R^2^ within a sphere of radius R projected onto a 2D plane (**Fig. 4A**) (28). This prediction matches simulations of hemozoin in a FV-like environment but does not account for hemozoin motion in expanded FVs where the TAMSD levels off at values that are much smaller than expected (**Fig. 3C**). This discrepancy suggests that physical micro-confinements within the FV limit crystal motion to sub-compartments of ∼ 1.1 μm radius that are 2-fold smaller than the expanded FV radius (∼ 2.5 μm) (***SI Appendix,* Tables S6, S8**). These additional barriers may be due to membranous remnants of vesicles that deliver host hemoglobin to the FV or other heterogeneities in the FV matrix that confine crystals (29).

**Figure 4:**
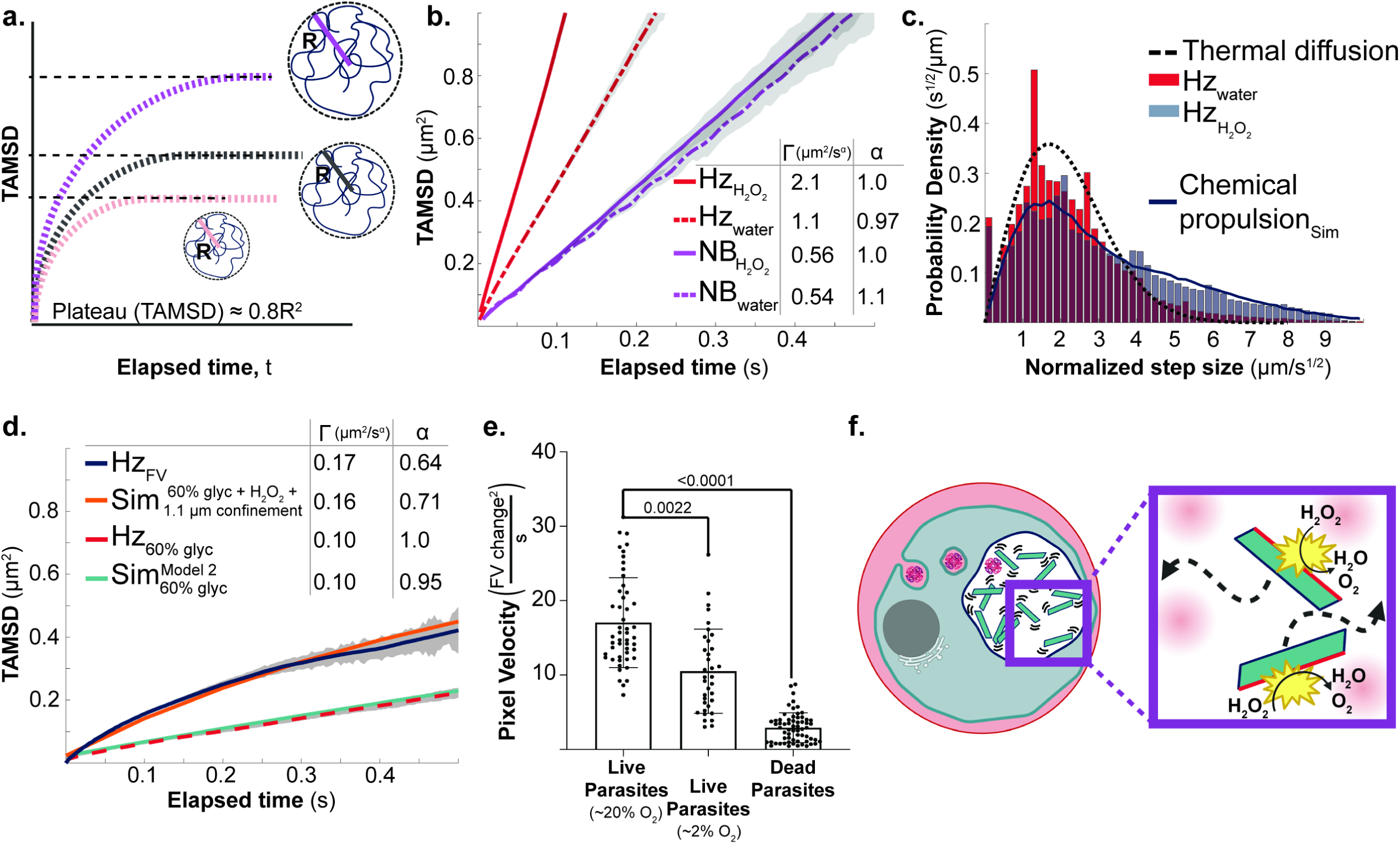
Extra energy and smaller confinement are required to capture crystal trajectories in the FV. **a**, Diagram showing predicted TAMSD plots for variable confinement radii. **b**, TAMSD of hemozoin crystals and inert nanobeads in water versus 15% aqueous H_2_O_2_. Error bars represent the standard deviation of the means of individual trajectories and are shown in grey. See Table S1 for fitted TAMSD values. **c**, Histograms showing probability density of average step size of each crystal over normalized time. Maroon bins show overlapping data in probability distributions. Solid curve is the step-size distribution predicted by our model of propulsive activity due to H_2_O_2_ catalysis. **d**, TAMSD of hemozoin crystals in expanded FVs compared to particles simulated with Brownian motion trajectories, experimental interaction potentials, confinement derivation from diagram in Fig. 4A and distribution of step sizes from Fig. 4C. See Table S8 for fitted TAMSD values. Error bars represent the standard deviation of the means of individual trajectories and are shown in grey. **e**, Population analysis of time-averaged mean squared pixel intensity change (TAMSI) of a FV in the specified conditions. Statistical analysis was done by Student’s t-test to determine p-values. **f**, Scheme depicting both stimulation and micro-confinement to explain hemozoin crystal trajectories in the FV. Inset: Pink compartments depict possible vesicle remnants within the FV matrix that confine hemozoin motion.

The enhanced diffusivity and heavy tails of longer step sizes of FV hemozoin suggested the possibility of energy inputs beyond thermal energy driving crystal motion (**Fig. 2D**). We first ruled out that white light of variable brightness affected hemozoin motion, as changing microscope light intensity from 1.8 to 25.3 mW/cm^2^ did not alter the anomalous exponent or diffusion coefficients in expanded vacuoles (***SI Appendix,* Fig. S5A**). We next tested a potential chemical origin for the enhanced diffusivity of FV hemozoin. Hydrogen peroxide (H_2_O_2_) is produced within the aerobic FV of parasites as a byproduct of hemoglobin digestion. This catabolic process releases ferrous heme, which oxidizes to form hemozoin and generates hydrogen peroxide from ambient O_2_ (30, 31). Malaria parasites do not encode catalase to degrade FV hydrogen peroxide, and how they neutralize this oxidative and cytotoxic stress remains unclear. Iron and other metals have been shown to catalyze hydrogen peroxide decomposition to release kinetic energy (32-36). We hypothesized that catalysis of this reaction within the FV by surface-exposed metals on hemozoin stimulates crystal motion.

We tested this hypothesis by first asking if hemozoin catalyzed H_2_O_2_ decomposition. We utilized an in vitro activity assay that used a surfactant to trap and visualize the O_2_ gas bubbles generated by this reaction (37). Both parasite-derived and synthetic hemozoin (β-hematin) but not inert nanobeads produced visible foaming, but bubbling by parasite hemozoin was notably stronger than synthetic crystals prepared from commercially sourced heme (***SI Appendix,* Fig. S6**). This result supports the conclusion that both parasite and synthetic hemozoin catalyze H_2_O_2_ decomposition but that parasite hemozoin has higher activity.

We next tested whether 15% H_2_O_2_ enhanced the diffusivity of isolated parasite hemozoin crystals in water. Using SPT, we observed a nearly 2-fold increase in the diffusion coefficient, which is comparable to previously reported motion enhancements for metallic nanoparticles based on this reaction (**Fig. 4B, *SI Appendix,* Fig. S5B, Table S1**) (38, 39). A milder enhancement was observed for synthetic hemozoin crystals, but this “boost” was not observed for inert nanobeads in water (**Fig. 4B, *SI Appendix,* Fig. S5B, Fig. S7, Tables S1, S3**). These motion enhancements matched the relative reactivities of H_2_O_2_ with parasite hemozoin, synthetic hemozoin, or inert nanobeads observed by the in vitro activity assay (***SI Appendix*, Fig. S6**). Power-law analysis of parasite hemozoin motion showed that 15% H_2_O_2_ did not substantially alter the α exponent from its value in water (α ∼ 1) indicating enhanced diffusion (**Fig. 2C, *SI Appendix,* Table S1**). However, the step-size histogram for isolated hemozoin crystals in 15% H_2_O_2_ displayed a heavy tail of longer step sizes, surprisingly similar to the histogram of FV hemozoin but in contrast to isolated parasite hemozoin in 60% glycerol (**Fig. 2D**, **Fig. 4C**). The experimental step-size histogram for isolated hemozoin in water (1 cP) was accurately fit to a single population with a diffusion coefficient of 1.44 μm^2^/s at a particle radius of 0.15 µm. In 15% H_2_O_2_, the histogram of crystal step sizes could only be accurately fit with a bimodal model where 38% of the crystal population had D_1_ ∼ 0.9 μm^2^/s and the other 62% of crystal step sizes fit a D_2_ ∼ 5.8 μm^2^/s (***SI Appendix,* Fig. S5C**). The close similarity between D_1_ and the thermal diffusion coefficient expected for 0.2 µm radius particles (1.1 μm^2^/s) suggests a smaller population of purely diffusive crystals and a larger population of crystals whose ≥5-fold larger D_2_ value resembles the enhancement ratio observed for FV hemozoin (***SI Appendix,* Table S5, Fig. S5C**).

We next investigated whether the crystal population with enhanced diffusivity could be explained by a simple model of random propulsion due to hemozoin surface catalysis of H_2_O_2_ decomposition. As a minimal model, we simulated 0.2 µm radius spherical particles undergoing thermal diffusion (D ∼ 1.1 μm^2^/s in water) and subject to randomized propulsive velocities normally distributed around zero. For translational propulsive velocity, we found that choosing a standard deviation of 60 µm/s for each direction (x, y, z) produced a step-size distribution that matched the faster fraction in the bimodal fit. Combining the step-size distribution of this active population (62%) with that of a thermally diffusing population (D_1_ ∼ 0.9 μm^2^/s, 38%) produced a step-size distribution like that observed experimentally (**Fig. 4C, *SI Appendix,* Fig. S5D**). Thus, our model for active propulsion of hemozoin by random catalysis of peroxide decomposition is sufficient to explain the experimental bimodal step-size distribution.

We next tested whether simulated particles in the estimated FV viscosity, with a smaller radial boundary of 1.1 µm and a bimodal distribution of step sizes parametrized from our hemozoin results in 15% H_2_O_2_, could quantitatively account for the enhanced diffusion and anomalous constraints of FV hemozoin. Inclusion of either the 1.1 µm radial boundary or the bimodal step-size distribution in simulations failed to match experiments, but including both properties nearly quantitatively matched the experimental TAMSD (**Fig. 4D**). The strong agreement between simulations and experiments supports our hypothesis that sub-compartmental constraints and chemical stimulation quantitatively explain the enhanced and anomalous features of FV crystal motion.

Our model predicts that a reduction in H_2_O_2_ concentration will suppress crystal motion. Since hydrogen peroxide generation in the parasite FV is expected to depend on delivery of oxyhemoglobin to this compartment by ingestion of RBC hemoglobin (31)(40), we reasoned that lowering ambient oxygen tension would diminish oxyhemoglobin levels and reduce H_2_O_2_ concentration in the FV. To test the effect of this reduction on crystal motion, we determined pixel velocities (TAMSI) for hemozoin moving within parasites grown and maintained at ambient 20% O_2_ or at reduced O_2_ tension of 2% expected to lower oxyhemoglobin levels by 80% (***SI Appendix,* Fig. S5E**). Critically, parasites have a low respiratory requirement for O_2_ and grow similarly in these two conditions (41). In contrast to unchanged parasite growth, there was a ∼2-fold reduction in pixel velocity for hemozoin motion in live parasites in 2% O_2_ that remained faster than observed for dead parasites (**Fig. 4E**). This decrease in speed directly supports our model that H_2_O_2_ serves as a fuel for propelling crystal motion in the FV and that lowering peroxide concentration slows hemozoin dynamics.

## Discussion

Hemozoin formation is both an essential survival mechanism and critical axis of drug activity against blood-stage malaria parasites. Although major structural properties of hemozoin are well studied, the rapid tumbling of these crystals within the parasite FV has remained a major enigma and defied experimental dissection. Our study incisively unravels the origin of this motion, which is both faster than diffusion and subject to surprisingly tight physical confinement (**Fig. 4F**).

Quantitative analysis of hemozoin movement provides a unique probe that has revealed structural obstacles within the FV matrix that constrain crystal displacements to regions that are smaller than FV boundaries. We hypothesize that these constraints may reflect membrane remnants of hemoglobin-uptake vesicles or other physical heterogeneities within the FV milieu similar to recent biophysical observations in yeast cytoplasm (29, 42). While the static nature of the *P. falciparum* FV has been visualized extensively by electron microscopy, the dynamic biophysical properties of this environment have remained inaccessible. Our analysis of hemozoin motion provides a rigorous estimate of 100-200 cP for the effective viscosity of the FV matrix that is similar to a mammalian lysosome. We anticipate that the experimental and computational approaches developed in this study will enable future studies of the FV to further test and refine understanding of this critical compartment and its key biochemical functions relevant for parasite physiology and antimalarial drug activity.

Hemozoin has been viewed canonically as a chemically inert detoxification product within the parasite FV (43, 44). Our results reframe the physiological role of these crystals and support the biological paradigm that metal-catalyzed H_2_O_2_ decomposition on the surfaces of hemozoin generates a random propulsive force to boost crystals towards larger steps that enhance diffusion while maintaining an unpredictable trajectory. This propulsive chemical mechanism is well-documented for synthetic metallic nanomachines, but *Plasmodium* hemozoin provides an unprecedented biochemical example of endogenous nanoparticle self-propulsion in cells (38, 39). Additional biological phenomena with this mechanism are likely to exist.

*Plasmodium* parasites do not encode a catalase enzyme, and mechanisms to neutralize the oxidative and cytotoxic stress of H_2_O_2_ production in the FV due to hemoglobin catabolism have been uncertain. Prior studies suggest possible roles for RBC-derived human catalase as well as parasite glutathione in reducing vacuolar H_2_O_2_ stress (30, 45). We propose that catalytic decomposition of peroxide on the surface of hemozoin also contributes substantially to H_2_O_2_ detoxification within the FV. Chemical strategies that interfere with hydrogen peroxide decomposition on the surface of hemozoin are therefore predicted to reduce crystal motion and increase oxidative stress for parasites that sensitizes them to therapeutic strategies targeting this vacuole. This hypothesis is consistent with prior works which reported that knockdown of a lipocalin-like protein in the FV diminished hemozoin tumbling and impaired parasite growth via oxidative stress that was reversed by the antioxidant Trolox (9, 10). The mechanistic connection between this lipocalin-like protein and hemozoin motion remains undefined but may involve differential exposure of surface metals on hemozoin due to altered crystal morphology and/or surface interactions in the absence of lipocalin.

The surface chemistry of catalytic H_2_O_2_ decomposition by metallic nanoparticles is complex (46), but our results provide initial insights into surface features of hemozoin that underpin this reaction in the FV. Hemozoin contains pentacoordinate iron within heme molecules that may be accessible at crystal surfaces and contribute to weak catalytic breakdown of H_2_O_2_ (3, 47-49). Differences in average crystal dimensions and surface area between synthetic and parasite-derived hemozoin may contribute to their apparent reactivity differences with H_2_O_2_. Parasite-derived hemozoin also has additional surface complexity and contains adhered lipids, proteins, and metals whose abundance and detection can vary with purification method and stringency (50). The higher apparent reactivity of H_2_O_2_ with parasite hemozoin than synthetic hemozoin may suggest that additional metals on the surface of parasite-derived crystals contribute to its stronger activity (***SI Appendix,* Fig. S6-S7**). Prior elemental analyses have detected calcium, silicon, and iron on the surface of parasite hemozoin (48, 51). Our previous work also suggests the presence of labile iron within the FV (52). Labile iron adhered to hemozoin crystals may underpin its enhanced activity with H_2_O_2_ compared to synthetic hemozoin. Surface-bound calcium or silicon, which do not catalyze H_2_O_2_ decomposition but enhance catalysis by other transition metals, may also contribute (53). The ability of metal chelators like EDTA and deferoxamine to quench catalytic H_2_O_2_ decomposition by parasite hemozoin supports a key role for surface labile metals (***SI Appendix*, Fig. S6**). We have ongoing interests and experiments to further understand hemozoin surface chemistry. It also remains an important future challenge to deeply understand the oxidative chemistry in the parasite FV that underpins H_2_O_2_ generation in this compartment.

Finally, dynamic motion of hemozoin within the FV matrix disperses crystals and maximizes their surface area available for biomineralization of labile heme released from hemoglobin proteolysis. Indeed, simple calculations suggest that crystal aggregation may reduce the exposed surface area up to 70%, and photodynamic ablation of hemozoin motion causes crystals to aggregate (***SI Appendix*, Fig. S8, Movies S7-S10**), as previously described (11). Multiple studies have also reported that isolated hemozoin crystals have an intrinsic tendency to clump after purification (54-56). These observations are consistent with the interaction potentials between crystals determined in this study from single particle tracking that indicated a shallow minimum at 0.82 µm, suggesting a weak attraction at the contact distance between crystals (**Fig. 3B**). Thus, hemozoin motion may serve an additional physiological role to maximize the exposed surface area of hemozoin crystals and to enhance the rate of heme detoxification within the FV, especially during peak stages of hemoglobin digestion.

## Materials and methods

***SI Appendix,* Figures S1-S9, Tables S1-S8, Movies S1-S10**

### Cell culture

#### Parasite culturing

*P. falciparum* Dd2 (42) and NF54 (57) parasites were grown in human red blood cells received from the University of Utah Hospital blood bank at 90% N_2_, 5% CO_2_, and 5% O_2_ at 37°C in RPMI 1640 containing AlbuMAXI (Thermo Fisher Scientific) supplemented with glutamine (2% hematocrit), as previously described (58). Medium was changed daily. Parasitemia was measured by Giemsa-stained blood smear or by flow cytometry (58). Parasites were routinely synchronized to the early ring stage by treatment with 5% D-sorbitol. All imaging of hemozoin motion was performed using late trophozoite- and schizont-stage parasites (36-48 hours post synchronization).

#### Parasite fractionation

Intact parasites within a permeabilized RBC ghost and isolated food vacuoles were prepared by membrane lysis. Parasitized red blood cells were permeabilized with 0.05% saponin in PBS, as described previously (58). This treatment released the hemoglobin contents in the RBC but maintained the membrane of isolated intact parasites. The plasma membrane of isolated parasites was permeabilized with exposure to ice-cold acidic hypotonic buffer (20 mM MES, pH 4.5), transferred to a watch glass on ice, mixed forcefully through a 27-G needle, and centrifuged at 15,800xg. The supernatant was removed by vacuum aspiration, and the pellet was resuspended in intracellular solution (120 mM KCl, 10 mM NaCl, 25 mM HEPES, 2 mM MgCl_2_, 5 mM Na_2_HPO_4_, pH 7.3) and triturated with a 27-G needle, as previously described (59). Isolated food vacuoles were resuspended on ice and imaged within 20 mins of isolation.

#### Inhibitor treatments

Parasites were magnet purified (isolating schizont stages) and incubated for 6 hours with fresh red blood cells. Newly invaded red blood cells were treated with 5% D-sorbitol (isolating ring stages) to produce rings with a 6-hour synchrony window. Parasites were separated into different wells and treated with high concentrations of inhibitors to induce cell death. Parasites were treated continuously with 0.2 μM WR99210 (Jacobus Pharmaceuticals) after ring synchronization (IC_50_: 0.043 μM); 0.1 μM chloroquine (Sigma-Aldrich) at 30 hours post-synchronization (IC_50_: 0.024 μM); 0.2 μM artemisinin (Sigma-Aldrich) at 30 hours post synchronization (IC_50_: 0.032 μM); or 200 μM 5-aminolevulinic acid (ALA, Sigma-Aldrich) at 24 hours post synchronization followed by 1 min of light exposure after 12 hours of ALA incubation (11, 60). Parasites were imaged at 36 hours post-synchronization and were scored based on moving vs. static hemozoin crystals. Thirty parasites were counted for each replicate and three replicates were taken for each drug treatment. Image analysis and TAMSI quantification was performed on chloroquine-treated parasites, although similar results were obtained for all drug-treatment conditions.

#### Synthetic hemozoin

Beta-hematin (synthetic hemozoin, sHz) was synthesized by adding 47.5 mg hemin (Sigma-Aldrich, H9039-1G) into 10 mL of 0.1N NaOH. Heme was precipitated slowly by adding 3.5 mL of glacial acetic acid dropwise. Tubes were incubated for 6 hours at 80°C and washed with 100 mM sodium bicarbonate, pH 9.1, as previously described (61).

#### Foaming Assay

Hydrogen peroxide decomposition leads to formation of water (H_2_O) and molecular oxygen (O_2_). In a microcentrifuge tube, 1% Triton X-100, water, and 15% H_2_O_2_ (final concentration) were added to 10 µL of packed pellet. Sample pellets included polystyrene nanobeads (NB), parasite hemozoin (pHz), synthetic hemozoin (sHz), parasite hemozoin with metal chelators (100 µM of deferoxamine (DFO, Sigma-Aldrich) or ethylenediaminetetraacetic acid (EDTA, Sigma-Aldrich)) and 100 units catalase (Sigma, C9332-1G). As a qualitative approach, trapped oxygen gas from this reaction was visualized as foam after 40 mins incubation, as previously described (37). Hydrogen peroxide concentration was confirmed by UV-vis absorbance spectroscopy (62).

### Microscopy

#### Widefield microscopy

Live parasites were imaged with a Nikon Ti-E inverted widefield microscope equipped with an Andor Zyla CMOS camera and 100X PlanAPO oil immersion objective. All pixel analysis time lapses were taken at ≥300 fps, while all single-particle tracking time lapses were taken ≥200 fps (slower frame rate due to larger region of interest). Microscope slides were incubated for 5 mins of “rest” after sample mounting and before imaging to minimize drift/bulk fluid flows under the microscope. All time lapse images were compiled as .nd2 files. Samples containing infected RBCs at 12-15% trophozoite or schizont parasitemia were mounted on a standard 75 x 25-mm glass microscope slide. For imaging, 7 μL of the resuspended culture was deposited on a slide and covered with a 22 x 40-mm coverslip. Even pressure was applied to the sample to remove excess liquid and the coverslip was sealed with clear nail polish to prevent evaporation. Low oxygen condition samples were imaged with flat-bottom dishes (CellTreat, #229632) on the same microscope, to ensure a sealed incubator that maintained low oxygen concentrations

#### Low O_2_ environment

An OKOLAB stage incubator on the Nikon Ti-E microscope was used with a dish adaptor. The clear lid was sealed with parafilm and flushed with CO_2_ for 20 minutes before images were collected. Samples were measured by an FD-103 Waterproof and Shockproof O_2_ Meter (Forensics Detectors) with a tube connection to the sealed incubator. Meter had range between 0-30% O_2_ with resolution within 0.1% O_2_ within < 1 minute.

#### Expanded vacuoles

Infected red blood cells were incubated in 100 μl of media diluted 1:1 with deionized water at 37°C for 20 mins before imaging. For imaging, 7 μL was deposited on a slide and sealed with a coverslip. Expanded vacuoles that retained hemozoin motion were imaged as described above.

#### Hemozoin isolation

A culture of infected RBCs at 1-20% parasitemia was saponin treated and resuspended in cold PBS. Parasites were spun down at 15,000xg for 5 mins, and resuspended in 2% SDS. Crystals were centrifuged at 15,000xg for 5 mins and resuspended in room temperature PBS with 2 μl of Qiagen protease and incubated for 1 hour at 37°C. Crystals were then washed with 2% SDS and spun at 15,000xg for 5 mins. Crystals were finally resuspended in 6M urea and stored in 4°C until imaging. Crystal pellets were washed twice and resuspended by vortexing in 100% water or 40-60% glycerol/water (w/v) concentrations for imaging.

#### H_2_O_2_ addition

30% H_2_O_2_ (Fisher, BP2633-500) was removed from a freshly opened bottle and mixed 1:1 with an equal volume of hemozoin crystals or nanobeads resuspended in water to give a final concentration of 15% H_2_O_2_ and immediately mounted onto a microscope slide. The viscosity of 30% H_2_O_2_ is 1.25 cP, which after mixing with water results in a final effective viscosity of 1.12 cP (63).

#### Light intensity

Light intensity on the Nikon Ti-E inverted widefield microscope was measured by a power meter sensor placed in the sample focal plane with a ThorLabs PM100A power meter analog console. Intensity was adjusted by with Sola automated LED lightsource.

#### Glycerol ratios and viscosity calculations

100% glycerol was added by weight converted to volume to distilled water to make aqueous glycerol solutions at room temperature (w/v). Solutions were vortexed for ≥ 1 hour to ensure adequate mixing. Viscosities were calculated from the simple Stokes-Einstein equation:

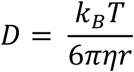

Where D is the diffusion coefficient derived from temperature (T), Boltzmann constant (k_B_), viscosity of surrounding medium (*η*) and the hydrodynamic radius of the particle (r). Viscosity values were solved by single-particle tracking control nanobeads with defined radius 0.4 μm at room temperature (22°C) and then computing and fitting the time-averaged mean squared displacement (TAMSD):

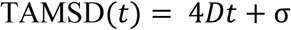

Where t is elapsed time. This model accounts for static localization error inherent to SPT data by introducing constant factor *σ* (27). Experimentally determined viscosities for nanobeads in the respective glycerol concentrations were nearly identical to literature values for these conditions (64). The effective viscosity from hypotonic expansion of vacuoles was estimated by using a weighted average:

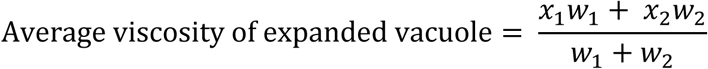

Where x is the estimated or calculated viscosity and w is percent of total expanded volume with that expected viscosity. We estimated wild type food vacuoles had viscosities similar to lysosomes (100 – 400 cP) and that the increase in volume from swelling was due solely to the influx of water (1 cP). We estimated that 90% of the swollen volume was water with 10% of the final volume having the original 100 cP lysosome viscosity, creating an effective 10 cP viscosity for the expanded vacuole.

#### Photodynamic ablation of hemozoin motion in parasites

Infected red blood cells were photosensitized by adding 200 µM exogenous 5-aminolevulinic acid (ALA) and incubated for 18 hours. This treatment results in an accumulation of the phototoxic intermediate protoporphyrin IX (PPIX), as previously published (11). Live parasites were imaged with a Nikon Ti-E inverted widefield microscope as described above, before and after light activation. Parasites were subjected to light activation by 1 min exposure to ∼450 nm light.

### Image analysis

#### Time-averaged mean squared pixel change intensity (TAMSI)

Single-particle tracking (SPT) proved challenging due to crystal congestion and overlapping trajectories within the small volume of the FV, limiting tracking to only a few frames. To overcome these limitations, we developed an alternative metric to assess crystal mobility: the time-averaged mean squared pixel intensity change (TAMSI) within the FV area as a function of elapsed time. Beyond approximately 0.2 s, TAMSI reached a plateau, indicating that crystal movement had fully decorrelated and longer elapsed times no longer contributed to increased pixel intensity changes. We empirically fit this time-dependent change to a power law growth function to determine the average characteristic time at which 50% of the plateau value was reached (see *Fitting* Methods and ***SI Appendix,* Fig. S9**):

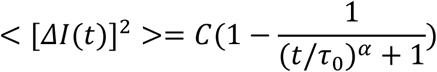

This analysis allowed us to define an apparent pixel velocity by dividing the TAMSI by the average characteristic time of ∼0.18 s, which we observed in native, living parasites. To compare between different FV-imaging conditions, we used the ∼0.18 s characteristic time to determine the pixel velocity for all time lapse experiments. Datasets were saved as .nd2 files and processed through MATLAB version R2021a with the Bio-Formats toolbox (65). Each frame of a dataset was adjusted so that bottom and top 1% pixel intensities were saturated to create better image contrast. The dataset was flipped to return the complement pixel intensities of each frame and the region of interest was defined manually by tracing the outer membrane of the food vacuole discernable in brightfield images. Each pixel intensity change was calculated by:

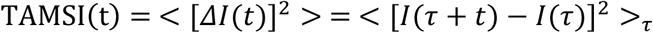

Where *I*(*t*) stands for pixel intensity, and the angle brackets denote averaging over various *τ* values and over all pixels. Each squared change was averaged over every pixel in the region of interest for every elapsed time *τ* and plotted as a function of increasing elapsed time.

#### Fitting

We observed nonexponential relaxation of pixel intensity change as a function of elapsed time. We used a general power law in the following form:

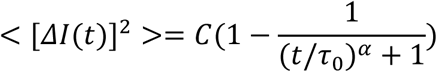

to empirically fit to the experimental data. The relaxation time *τ*_0_ encodes the time t at which pixel intensity change reaches half of its asymptotic value C to which the function tends in the limit of *t* → ∞. We used `scipy.optimize.curve_fit` function in Python to empirically fit our data (66). For dead parasites, the dominant contribution to pixel intensity fluctuation was the background noise that deviated from the power law equation used for conditions with moving hemozoin. Since we observed noise as the dominant feature, we instead used only the first term of expansion of the power law in Poiseux series (for *α* = 0.5, *C* = 0.05):

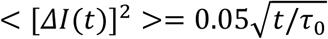

Before fitting, we subtracted from the experimental data the short-time average computed between 5th and 15th data points. Then, we performed fitting from the 10th data point onwards, where we observed an extremely slow increase. As the number of particles in a small pixel is governed by Poisson distribution, it depends only on the total intensity divided by the number of pixels. For long times *t* the Poisson random variables are going to be statistically independent, and in principle *C* can be predicted from the Skellam distribution properties (67). Thus, we expected *C* to be comparable between dead and living parasites. Thus, the choice of *α* and *C* were based on the median values observed for the infected red blood cell group.

### Single-particle tracking

#### TrackMate

We used the Fiji plugin TrackMate to single-particle track hemozoin crystals throughout time lapse images (68, 69). We chose the LoG (Laplacian of a Gaussian) detector as our method of localization. This method applies a LoG filter to each image to find local maxima with quadratic fitting for sub-pixel localization. We estimated our object diameter to be 0.6 μm unless tracking the defined 0.8 μm diameter nanobeads. Each time lapse was manually processed by resetting the initial thresholding to rule out background spots. We used the Simple LAP tracker because we assumed there would be no branching tracks (particles were not breaking and reforming). We used 0.5 μm as our linking max distance, 0.5 μm for our gap-closing max distance, and 2 as our gap-closing max frame gap. Tracks were minimally filtered by particle spots that were “stuck” where their displacements did not increase with time.

#### Time-averaged mean squared displacement

For each system, we recorded numerous tracks of different length (number of time points), and different time resolution (differing from experiment to experiment). To analyze such heterogeneous dataset, we first computed and stored all displacements, i.e., *r*(*i* + *j*) − *r*(*j*); *i*, *j* ∈ ℕ, and used them as a basis for the subsequent analyses. To compute time-averaged mean squared displacements (TAMSD), we iterated over the list of displacements, squared them, and averaged ones belonging to equal lag i. To i=1, we attributed a timestep averaged over all experiments belonging to the same system. To compute uncertainties, we repeated the computation of squared displacements (SD) but limiting the computation only to those displacements which are statistically independent. For instance, for the same particle, r(3)-r(0) is not statistically independent from r(5)-r(2). Therefore, we used only these SDs for which start time *j* is a multiple of the lag time i. For each lag time i, we stored a list of N_i_ statistically independent SDs. Subsequently, for each lag time i, we performed 100 random selections of N_i_ SD values from the list and used standard deviation of the means of the selected samples as a measure of uncertainty in TAMSD value. If there was only 1 value in the SD list, we attributed infinite uncertainty.

We fit TAMSD with two diffusion models:

Normal:

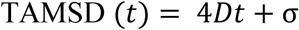

And anomalous:

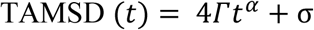

We used the `scipy.optimize.curve_fit` function in `python` to empirically fit the data and provided the uncertainties as inverses of fitting weights. This means that in least squares optimization, the residuals are divided by the uncertainties, making the points with poor statistics weigh less. We fit only the short-time parts of the TAMSD curves (0.1-0.4 s). To ensure that using averaged timesteps did not distort the results, for the systems in which normal diffusion is expected, we divided all the squared displacements by the timestep:

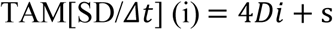

This procedure led to very similar values (*SI Appendix,* Tables S2 and S4 compared to Tables S1 and S3, respectively).

### Simulations

#### Control simulations for TAMSI comparison

We used 1/*τ*_0_ as a proxy for the diffusion coefficient. To test the accuracy of this approximated value, we performed a series of point-particle Brownian dynamics simulations in which we computed pixel intensity change using the same protocol as when analyzing experimental data, but instead of computing intensity change we computed the number of particles in each pixel. We observed faster relaxation (smaller *τ*_0_) for particles with higher diffusion coefficients *D*. Specifically, we performed Brownian dynamics simulations of 600 noninteracting particles of diffusion coefficients D = 2, 10, 20 μm^2^/s in a 20 nm × 20 nm × 20 nm box (with periodic boundary conditions) for 10000 steps (dt = 500 ps). Then, we divided the system into 100 2D pixels along X and Y dimensions and quantified the number of particles N in each pixel in time. We calculated:

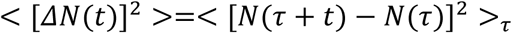

and averaged over the pixels. In agreement with Skellam distribution properties, the plateau occurred at twice the average number of particles in a single pixel, which is 2 × 600 / 100 = 12; *τ*_0_ is the value of time at which pixel particle number fluctuation reaches a value of 6. We observed that *τ*_0_ decreases with increase in diffusion coefficient, making its inverse a valid proxy for diffusivity. See Fig. S2d and Fig. S9 for results. A similar method was used recently employing a “countoscope” analysis (70). We noted that the simulations are scale invariant, so rescaling the length unit (in pixel size and diffusion coefficients) leads to the same results.

#### Brownian dynamics simulations

We performed Brownian dynamics simulations of hemozoin-like cuboidal “bricks” using a 1:1:3.5 ratio rectangular prism with dimensions 170x170x585 nm using custom MATLAB version R2021a code (54). Nodes were spaced ∼80 nm apart to total 74 nodes on the surface of each brick. These nodes had regularized stokeslets applied and summed over the entire brick. We calculated randomized rotational and translational steps based on the Stokes-Einstein equation for spherical particles in low Reynolds number. Timestep was 0.0033 s to match timesteps of experimental time lapses, temperature: 293.15 K. Confinement was realized by defining spherical region of various radii and defining repulsive Lennard-Jones interaction between wall and particles. Particle-particle interactions were defined by the same repulsive Lennard-Jones equation with epsilon estimated from published literature (*ε* = 0.08 kcal/mol, *σ* = 57 nm):(71)

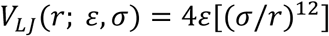

See Table S6 for physical parameters of each simulation, which mimicked experimental expanded FV conditions.

Using a custom MATLAB version R2021a code,(54) we modeled a sphere with radius 0.2 μm undergoing thermal translational and rotational diffusion in water due to Langevin forces. In addition, each sphere had propulsion due to chemical activity modeled by a fixed-body translational and rotational velocity. The propulsive velocities of the population of spheres obeyed normal distributions with mean zero. For rotational propulsive velocity, the normal distribution had a standard deviation of 314 rad/s for all three directional components (x, y, z), which is consistent with tumbling rotation speeds we observe in **Movie S4** (∼ 168 – 419 rad/s). For translational propulsive velocity, a standard deviation of 60 μm/s adequately reproduced the step-size distribution for the active portion of the experimental bimodal distribution for hemozoin in 15% H_2_O_2_. We simulated 1000 trajectories up to 0.33 s to mimic experimental conditions.

We performed Brownian dynamics simulations of 40 spheres in confinement using LAMMPS software (`fix brownian`) (72). Timestep was 1 μs, total number of steps was 20000000, and temperature: 293.15 K. Friction coefficients corresponding to 0%, 40%, and 60% glycerol were: 1917.0, 7131.0, and 20704.0 g mol^-1^fs^-1^. These friction coefficients translate to a hemozoin dynamic radius of 0.169 μm. Confinement was realized by defining spherical region of radius 2.9 μm and defining a short-range harmonic repulsive interaction between wall and particles (*ϵ* = 0.1kcal/mol/Å^2^, *σ* = 0.1μm). Simulations with “peroxide boost” were performed via custom `python` code, using the distribution in step sizes from experimental data and implementing hard spherical boundaries.

#### Parametrizing experimental crystal-crystal interaction

From tracks of hemozoin in the food vacuole, we computed histograms of distances between the separate crystals P(r). We used the Direct Boltzmann Inversion (DBI) method (73) to parametrize the potentials:

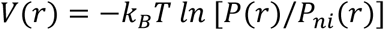

where P_ni_(r) is the histogram as if the crystals would not interact at all (ideal gas). We modeled the geometry underlying experimental data as a 3D sphere projected onto a 2D plane. Thus, to compute P_ni_(r) we performed the following computation:

1. Random-uniform insertion of 10000 point particles into 3D sphere of radius 2.9 μm
2. Projection of coordinates onto the XY plane.
3. Computing distances between all particle projections.
4. Computing histogram of distances, with 100 bins spanning a range from 0.0 to 6.0 μm.

We fit a 6-12 Lennard Jones potential:

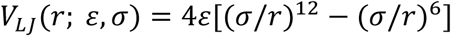

to the resultant DBI potential for distances larger than 0.64 μm (to avoid large errors stemming from fitting to a very steep increase due to excluded volume). Parameters giving best match were: ε = 0.24±0.06 kcal/mol, σ = 0.643±0.008 μm.

To show the robustness of our results to a specific form of the potential, we also performed simulations with two different interaction potentials. First, we used Weeks-Chandler-Andersen (WCA) potential — LJ-based exclusively repulsive potential representing volume exclusion:

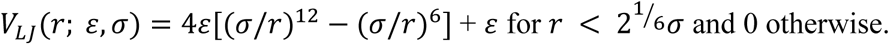

with ε and σ remaining the same as in DBI LJ potential. Second, we used Yukawa potential — exclusively repulsive potential with long-range repulsion:

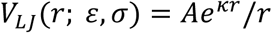

with A = 0.3 kcal/mol⋅μm, κ = 1 μm^-1^. Global cutoff was set to 2 μm.

#### Jump distribution

To extend the analysis, we computed histograms of the displacement lengths divided by 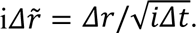 For a Brownian motion, such histograms are governed by the following probability density function:

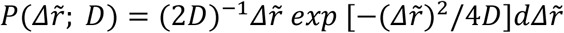

This function has a maximum for 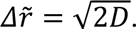 For low-viscosity systems, we computed histograms for i=1 to maximize the sample size. For high-viscosity systems, we computed histograms for i=10 to avoid contributions of static localization error to the jump statistics. Static localization errors are inherent to SPT data, especially in high viscosity systems with confinement, where the accuracy between the observable particle position is slightly different than the particle’s true position (27). We observed that fitting with the theoretical expression gives poor results, especially at high-jump length heavy tails. To quantify these tails, we performed fitting with bimodal model, i.e.:

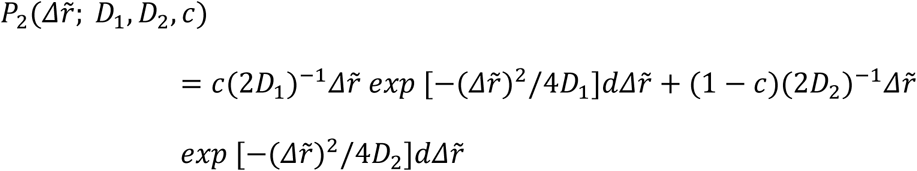

obtaining much better agreement with the experimental curves.

## Data availability

All data described in the manuscript are shown in the figures and provided in SI Appendix. Supplementary movies are provided in figshare. All time lapses for data from pixel analysis (TAMSI) in Fig. 1B-D and Fig. 4E are provided in figshare:

Movies - RBCs. figshare. Dataset. https://doi.org/10.6084/m9.figshare.28719737.v1

Movies - RBCs low O2. figshare. Dataset. https://doi.org/10.6084/m9.figshare.28719716.v1

Movies - Isolated parasites. figshare. Dataset. https://doi.org/10.6084/m9.figshare.28719722.v1

Movies - Isolated FVs. figshare. Dataset. https://doi.org/10.6084/m9.figshare.28719734.v1

Movies - Dead parasites. figshare. Dataset. https://doi.org/10.6084/m9.figshare.28719731.v1

Movies - Expanded vacuoles. figshare. Dataset. https://doi.org/10.6084/m9.figshare.28719719.v1

Supplementary Movies. figshare. Dataset. https://doi.org/10.6084/m9.figshare.28835870.v2

## Code availability

All MATLAB, python, and LAMMPS scripts used for data analysis and simulations, along with default parameters and all experimental single-particle tracks for step-size distribution and TAMSD analysis are available at https://github.com/emhastin/HemozoinMotion.git.

## Acknowledgements

We thank Kiarash Samsami, Ludivine Sanchez-Arias, and Sigala lab members for helpful discussions. We thank Xiang Wang, Mike Bridge, and Anton Classen at the Utah Cell Imaging Core Facility; Shawn Colby at the Health Sciences Machine Shop; and Brian Van Devener at the Utah Nanofab for experimental assistance. This work was supported by NIH grants R35GM133764 (PAS), R21AI185746 (PAS), and R35GM14749 (TCB); a pilot award from the Utah Center for Iron and Heme Disorders U54DK110858 (PAS); and a pilot award jointly funded by the Price College of Engineering and the 3i Initiative at the University of Utah (PAS, HF, and TCB). EMH was funded in part by NIH training grant T32AI055434. Brightfield microscopy and flow cytometry were performed using core facilities at the University of Utah.

