## Supporting Figures and Tables for "Chemical propulsion of hemozoin crystal motion in malaria parasites"

**This PDF file includes:**

Figures S1 to S9
Tables S1 to S8
Legends for Movies S1 to S10

### Figures S1-S9

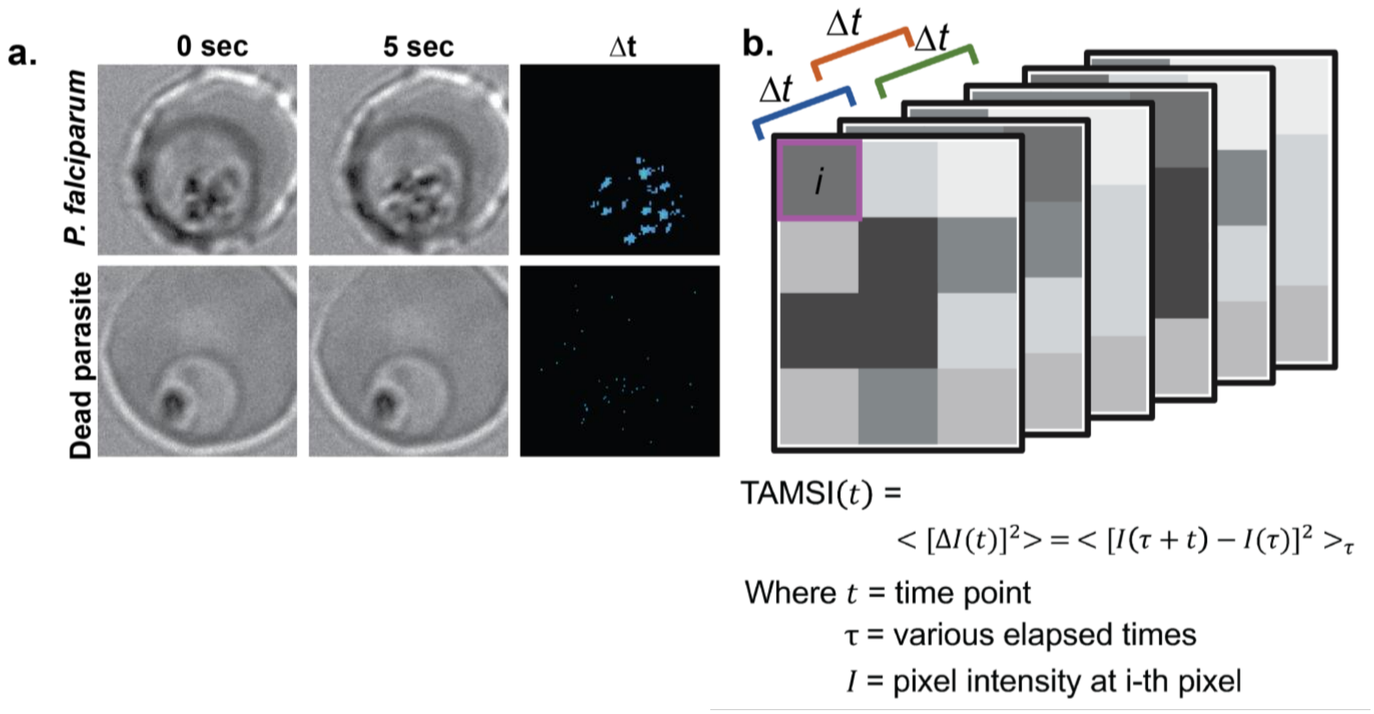


**Fig. S1: Time lapse image analysis of parasite hemozoin motion. a**, Live time lapse images of parasite-infected RBCs with pixel-by-pixel subtraction over 5 seconds and heat maps to display biggest pixel differences. **b**, Diagram explaining image analysis where each pixel's intensity is subtracted over consecutive frames and averaged within region of interest to define average squared change over elapsed time.

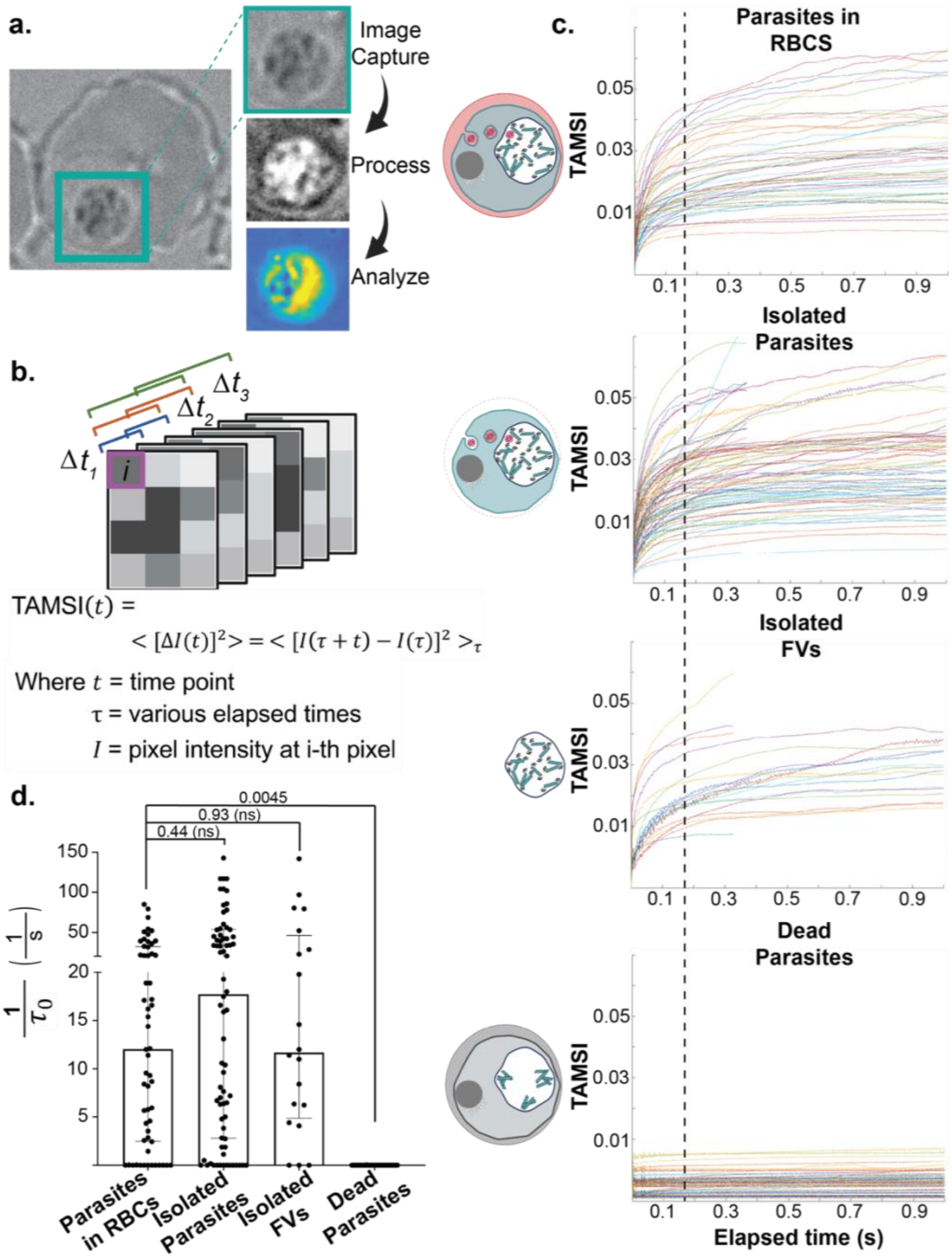


**Fig. S2: Fractionation and quantitative analysis of hemozoin motion. a**, Example of image analysis method where food vacuole is captured at above frame rate threshold, preprocessed to normalize pixel intensity values, then analyzed with overall pixel intensity change over time. **b**, Diagram explaining image analysis where each pixel's intensity is subtracted over different elapsed times, squared, and averaged within region of interest to define change in progressively longer elapsed times. **c**, Cumulative data from different time lapse experiments indicating a constant elapsed time (~0.18 s) was chosen to determine velocities of all conditions. **d**, A general power law fits experimental data in Fig. S2C where $1/\tau_0$ can be used as a proxy for diffusion coefficients. This correlation was corroborated by Brownian dynamics simulations where both TAMSI and diffusion coefficients were calculated for each data set and larger $1/\tau_0$ had a higher diffusion coefficient. See Fig. S9. Each point indicates $1/\tau_0$ of the same experimental data in Fig. 1D. For dead parasites, only the first term in expansion of the power law in Poiseux series was used. Statistical analysis was done by Student's t-test to determine the indicated p values.

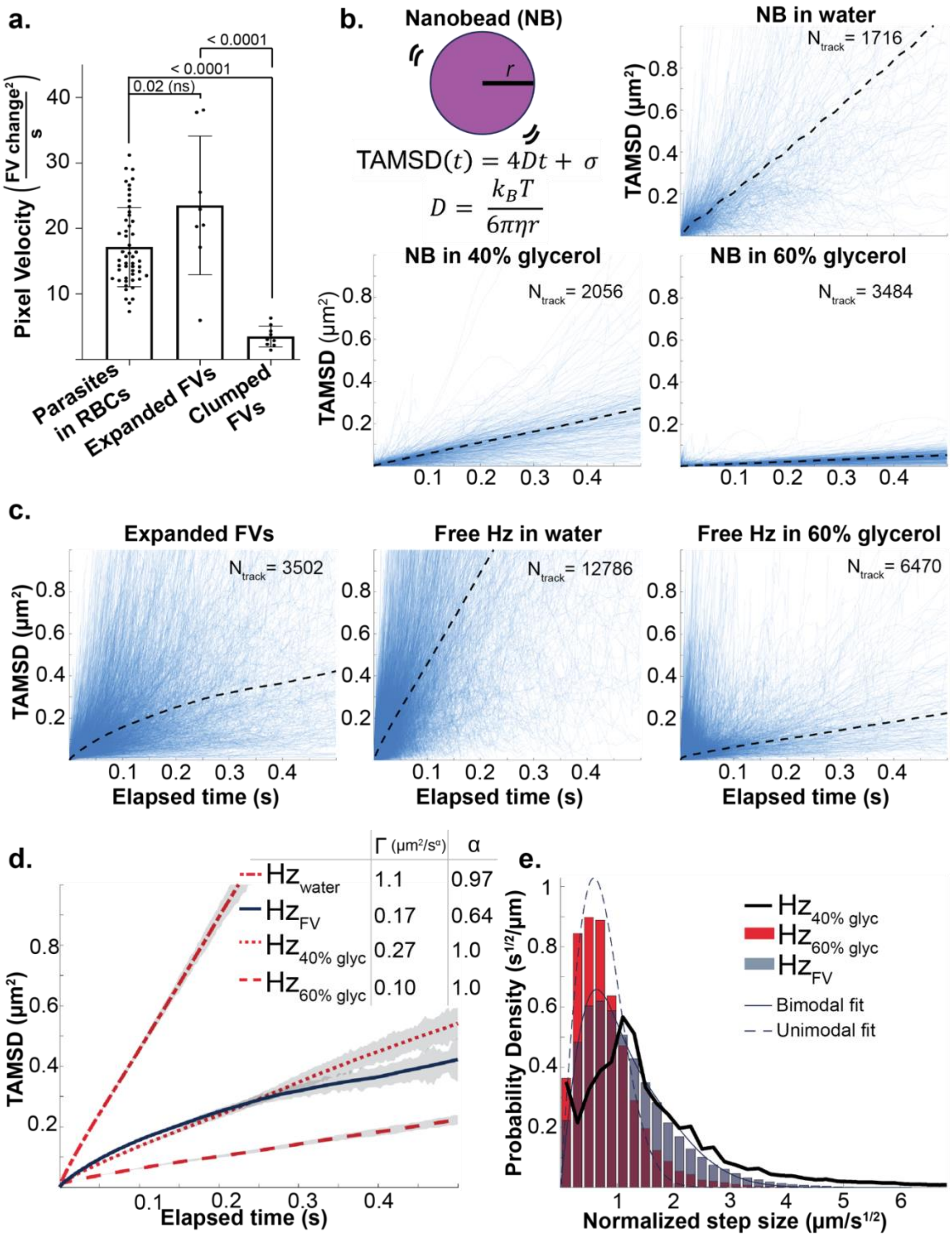


**Fig. S3: Single-particle tracking analysis of hemozoin motion. a**, Image analysis comparing different conditions using same TAMSI methods as Fig. 1D. **b**, Viscosities and SPT parameters from glycerol:water ratios were confirmed by Stokes-Einstein equation with 0.4 μm radius nanobeads (NB) including a static localization error in single-particle tracking (σ). Blue lines indicate individual track time-averaged mean squared displacements (TAMSDs) with dashed black line showing weighted average. **c**, Blue lines indicate individual track TAMSDs with dashed black line showing weighted average. **d**, TAMSD. Error bars represent the standard deviation of the means of individual trajectories and are shown in grey. N indicates the number of tracks. **e**, Probability density distribution of isolated crystals in 40% glycerol. Experimental step sizes showing difference between unimodal and bimodal fits.

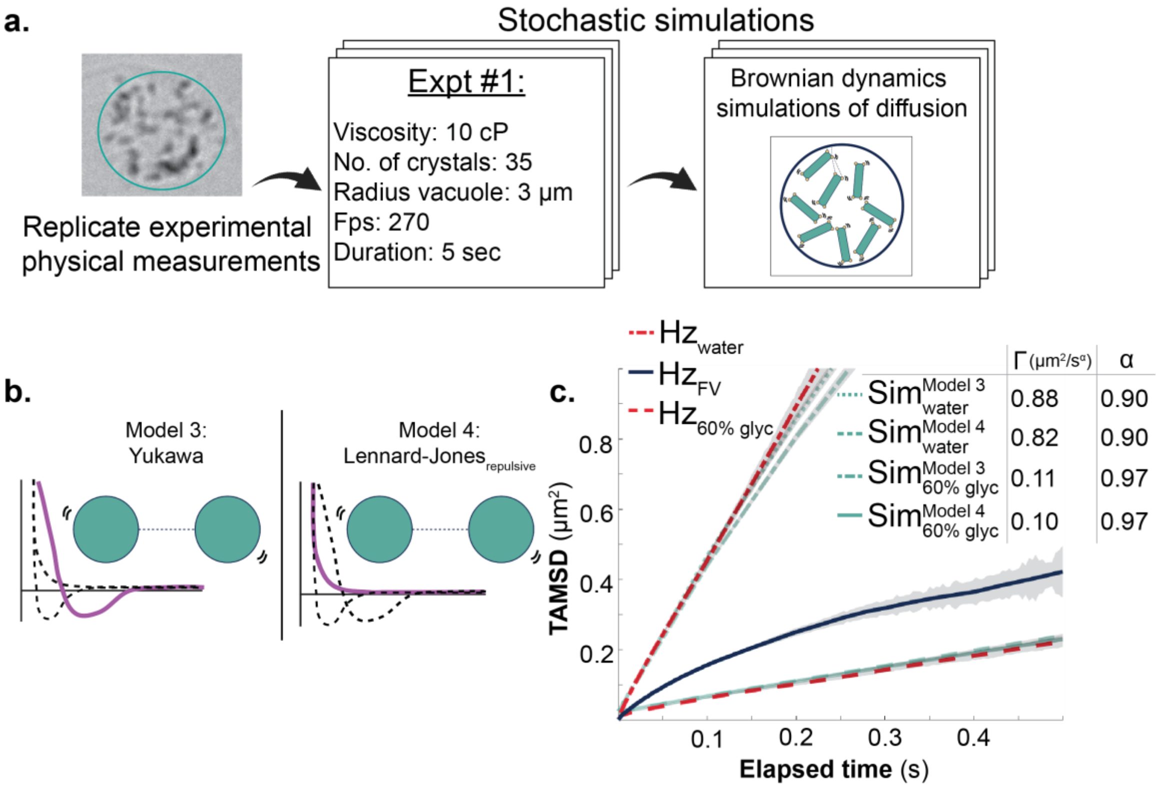


**Fig. S4: Brownian dynamics simulations of hemozoin motion to test boundary effects and particle-particle interactions**. **a**, Diagram showing example physical parameters that were used in each simulation, see Table S6. The FV radius, expected viscosity, number of crystals, and frame rate of each experimental expanded FV was modeled. **b**, Additional particle-particle interactions were evaluated including the Yukawa potential and the repulsive term of Lennard-Jones applied to spheres. **c**, TAMSD of hemozoin crystals in expanded FVs compared to particles simulated with Brownian motion trajectories and interaction potentials using Models 3 and 4 from B. Error bars represent the standard deviation of the means of individual trajectories and are shown in grey.

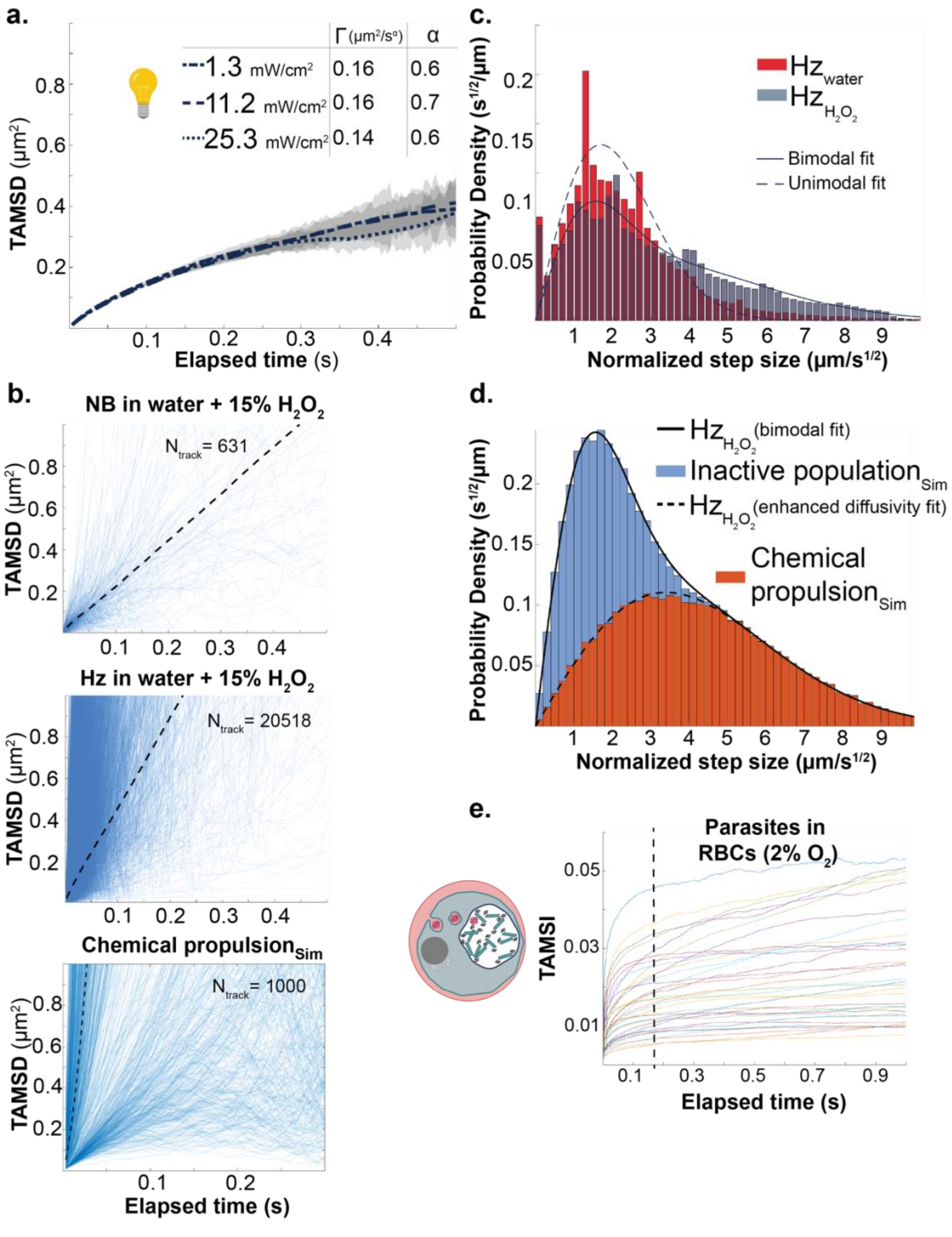


**Fig. S5: Impact of light and hydrogen peroxide on hemozoin motion. a**, TAMSD of hemozoin crystals in expanded FVs under different measured light intensities. Error bars represent the standard deviation of the means of individual trajectories and are shown in grey. **b**, Blue lines indicate individual track TAMSDs with dashed black line showing weighted average. Lowest TAMSD shows simulated tracks from chemical propulsion model. N indicates the number of tracks. **c,** Experimental step sizes showing difference between unimodal and bimodal fits. **d,** Histogram of step-size distribution simulated using a propulsion model. Step sizes of active population (orange) closely fit the expected step sizes for a particle with a diffusion coefficient of 5.8 $\mu m^2/s$ (dashed line). Adding the step sizes of the purely diffusive population (blue) shows the total distribution closely matches the bimodal fit for hemozoin in 15% peroxide from Table S5 (solid line). **e**, Cumulative data from different time lapse experiments indicating a constant elapsed time (~0.18 s) was chosen to determine velocities of all conditions.

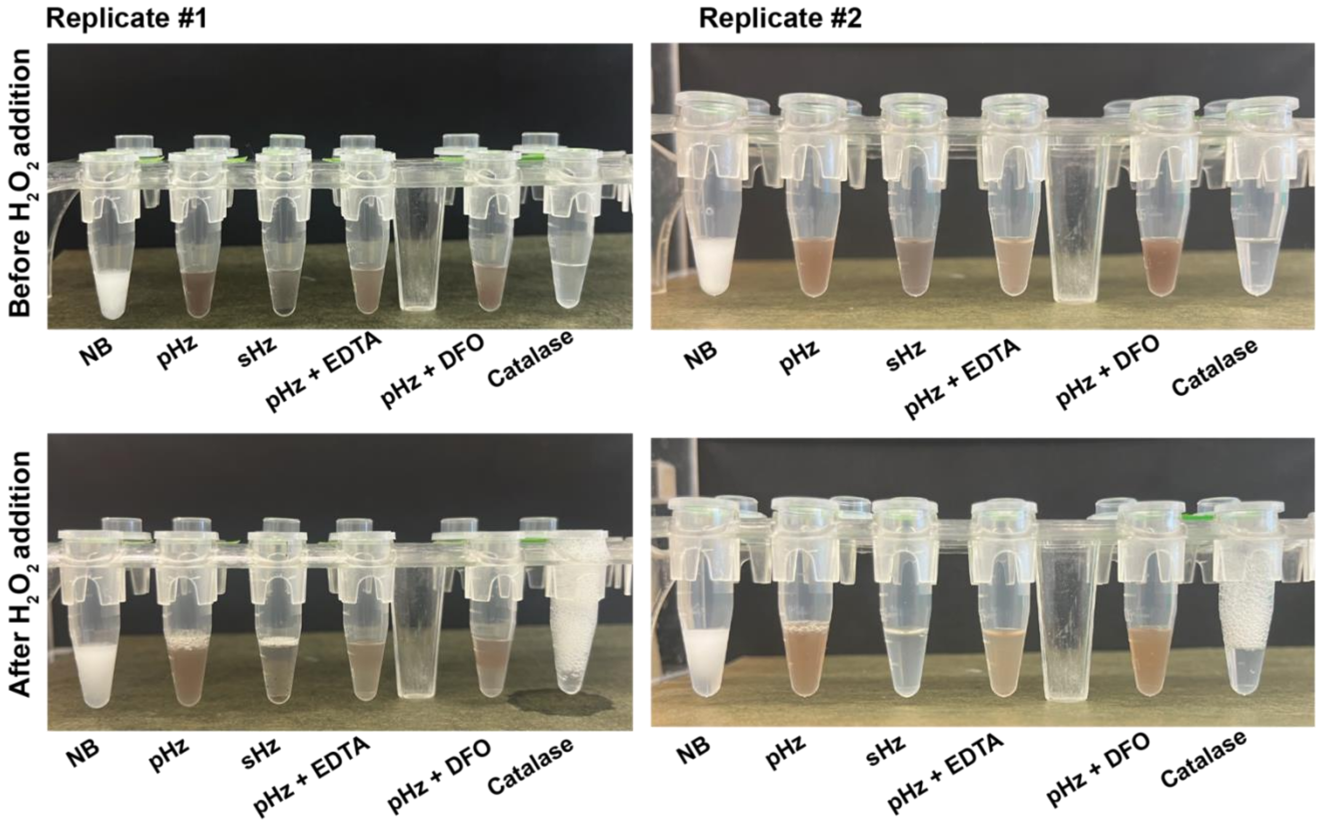


**Fig. S6: Foaming assay to test catalytic peroxide decomposition and inhibition by metal chelators.** Two independent replicates of foaming assay before and after addition of 15% $H_2O_2$. From left to right: 10 µL polystyrene nanobeads (NB), 10 µL resuspended parasite-derived hemozoin crystals (pHz), 10 µL resuspended synthetic hemozoin crystals (sHz), 10 µL parasite hemozoin with 100 µM EDTA (metal chelator), 10 µL parasite hemozoin with 100 µM DFO (iron chelator), 100 units catalase.

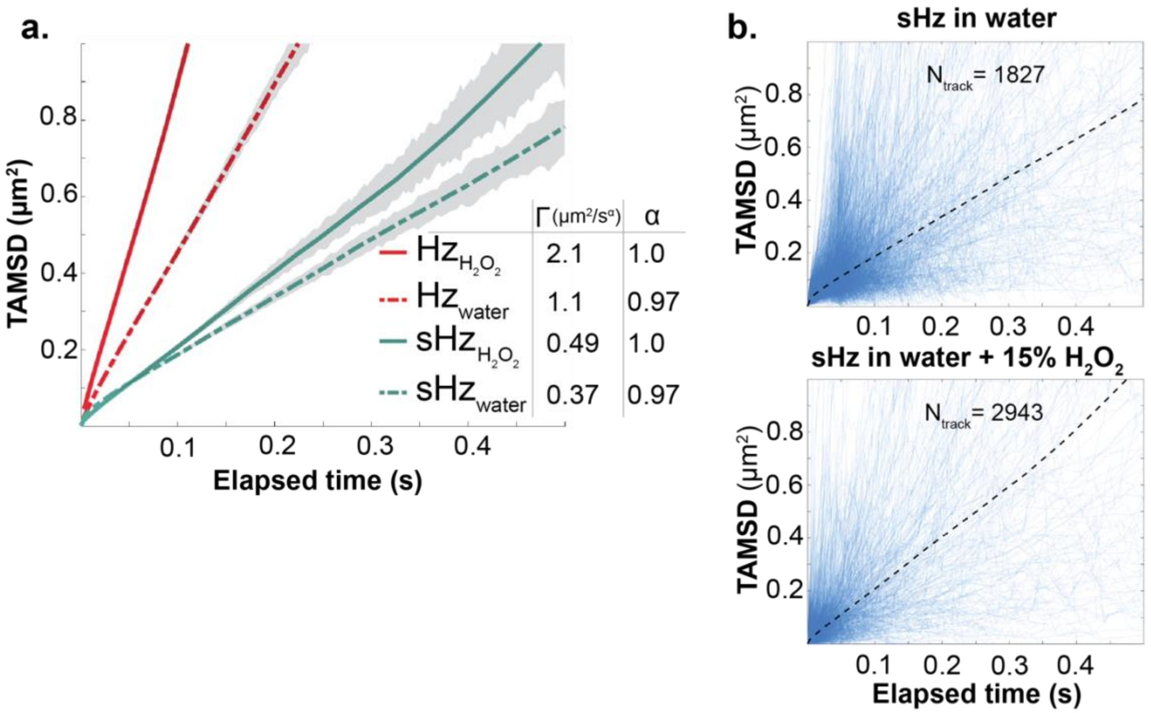


**Fig. S7: Single-particle tracking analysis of synthetic hemozoin motion. a,** TAMSD of synthetic hemozoin crystals (sHz) versus parasite-derived hemozoin (Hz) in water or 15% $H_2O_2$. Data for parasite-derived hemozoin is from Fig. 4. Error bars represent the standard deviation of the means of individual trajectories and are shown in grey. **b**, Blue lines indicate individual track TAMSDs with dashed black line showing weighted average. N indicates the number of tracks.

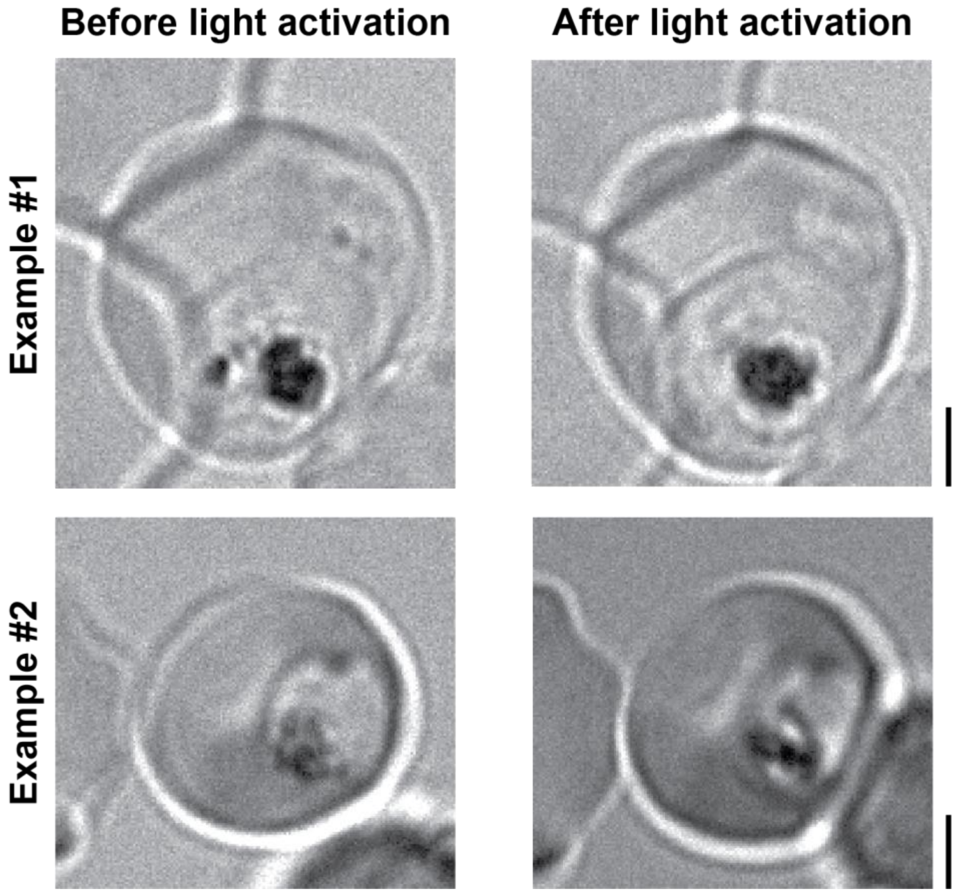


**Fig. S8: Hemozoin crystals clumping in the absence of dynamic tumbling.** Two examples of still images from time lapses showing the smaller and denser crystal area after photodynamic ablation of hemozoin motion. Example #1 can be seen in Movies S7-S8. Example #2 can be seen in Movies S9-S10. Parasites were cultured in 200 µM 5-aminolevulinic acid to induce photosensitizing porphyrin accumulation. Time lapse movies were recorded on the brightfield channel before and after illumination for 1 minute with ~450 nm light. Scale bars = 2 µm.

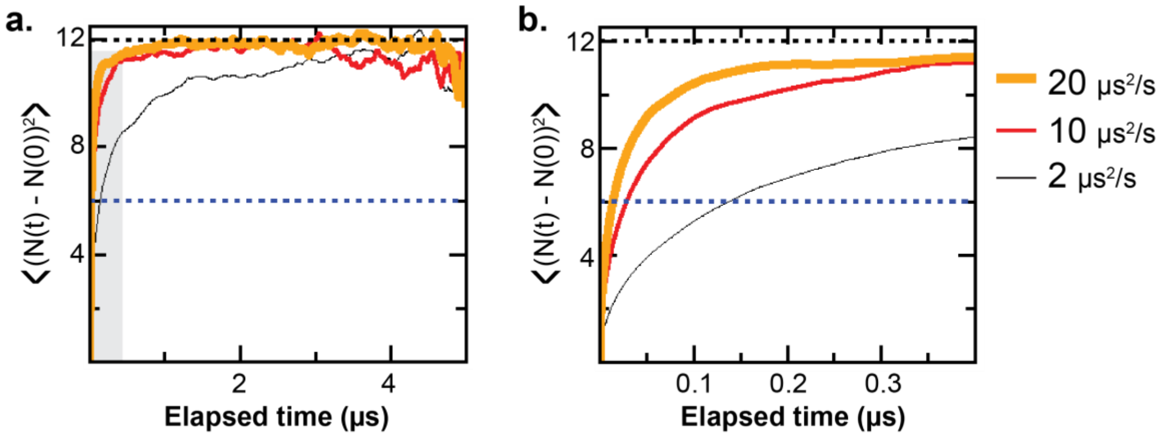


**Fig. S9: Relaxation time of time-averaged mean squared pixel intensity change serves as proxy of diffusion coefficient. a**, Dependence of time-averaged mean squared pixel intensity change (TAMSI) changes with diffusion coefficient obtained from Brownian dynamics simulations. **b**, Zoom into the region in which the curves cross the vertical line denoting half of the plateau value (blue).

**Supporting Tables S1-S8**

**Table S1.** Results of fitting anomalous and normal diffusion models to single-particle tracking TAMSD data with static localization error correction. Error bars based on curve fitting, as described in Materials and methods. For σ, values without parentheses indicate fit to the anomalous model ($4\Gamma t^{\alpha}+\sigma$) while values inside parentheses indicate fit to the linear model ($4Dt+\sigma$).

| | α | Γ (μm²/s^α) | D (μm²/s) | σ (μm²) |
|---|---|---|---|---|
| hemozoin in 15% peroxide (all) from 0.1 s to 0.4 s | 1.03±0.09 | 2.09±0.14 | 2.04±0.04 | 0.03±0.13 (0.00±0.02) |
| synthetic hemozoin in 15% peroxide (all) from 0.1 s to 0.4 s | 1.00±0.09 | 0.49±0.03 | 0.489±0.008 | 0.01±0.03 (0.011±0.005) |
| hemozoin in water (all) from 0.1 s to 0.4 s | 0.97±0.06 | 1.06±0.04 | 1.079±0.014 | 0.00±0.05 (0.024±0.009) |
| synthetic hemozoin in water (all) from 0.1 s to 0.4 s | 0.97±0.09 | 0.37±0.02 | 0.374±0.006 | 0.03±0.03 (0.037±0.004) |
| hemozoin in 40% glycerol (all) from 0.1 s to 0.4 s | 1.04±0.08 | 0.269±0.015 | 0.261±0.004 | 0.039±0.014 (0.031±0.004) |
| hemozoin in 60% glycerol (all) from 0.1 s to 0.4 s | 1.04±0.08 | 0.102±0.006 | 0.0991±0.0015 | 0.026±0.006 (0.0232±0.0011) |
| hemozoin in food vacuole (length≤300) from 0.1 s to 0.4 s | 0.64±0.12<br>1.8mW/cm²:<br>0.6±0.2<br>11.2mW/cm²:<br>0.7±0.2<br>25.3mW/cm²: | 0.173±0.004<br>1.8mW/cm²:<br>0.159±0.007<br>11.2mW/cm²:<br>0.164±0.007<br>25.3mW/cm²: | 0.200±0.004<br>1.8mW/cm²:<br>0.185±0.008<br>11.2mW/cm²:<br>0.189±0.006<br>25.3mW/cm²: | 0.00±0.04<br>(0.082±0.003)<br>1.8mW/cm²:<br>0.00±0.08<br>(0.083±0.006)<br>11.2mW/cm²: |

| | | | | |
|---|---|---|---|---|
| | 0.6±0.2 | 0.144±0.006 | 0.166±0.006 | 0.00±0.05 (0.071±0.004) 25.3mW/cm$^2$: 0.00±0.07 (0.087±0.004) |

**Table S2.** Results of fitting normal and anomalous diffusion models to single-particle tracking TAM[SD/$\Delta t$] data with static localization error correction, where i is time lag. Error bars based on curve fitting, as described in Materials and methods. For $\sigma$, values without parentheses indicate fit to the anomalous model ($4\Gamma i^{\alpha}+s$) while values inside parentheses indicate fit to the linear model (4Di+$s$).

| | α | Γ ($\mu m^2/s^{\alpha}$) | D ($\mu m^2/s$) | $s$ ($\mu m^2/s$) |
|---|---|---|---|---|
| hemozoin in 15% peroxide (all) from 30 to 140 | 1.00±0.07 | 2.0±0.8 | 1.96±0.03 | 10±30 (11±6) |
| hemozoin in water (all) from 30 to 140 | 1.02±0.05 | 1.0±0.2 | 1.117±0.012 | 10±12 (7±2) |
| hemozoin in 40% glycerol (all) from 30 to 140 | 0.99±0.06 | 0.27±0.09 | 0.262±0.004 | 11±4 (11.3±0.8) |
| hemozoin in 60% glycerol (all) from 30 to 140 | 1.01±0.06 | 0.10±0.03 | 0.1017±0.0014 | 8±2 (7.6±0.3) |

**Table S3.** Results of fitting anomalous and normal diffusion models to single-particle tracking TAMSD data with static localization error correction. Error bars based on curve fitting, as described in Materials and methods. For $\sigma$, values without parentheses indicate fit to the anomalous model ($4\Gamma t^{\alpha}+\sigma$) while values inside parentheses indicate fit to the linear model ($4Dt+\sigma$).

| | α | Γ ($\mu m^2/s^{\alpha}$) | D ($\mu m^2/s$) | $\sigma$ ($\mu m^2$) |
|---|---|---|---|---|
| nanobeads in peroxide (all) from 0.1 s to 0.4 s | 1.01±0.12 | 0.56±0.04 | 0.554±0.012 | 0.00±0.05 (0.000±0.008) |
| nanobeads in water (all) from 0.1 s to 0.4 s | 1.10±0.12 | 0.54±0.05 | 0.507±0.012 | 0.04±0.04 (0.007±0.008) |
| nanobeads in 40% glycerol (all) from 0.1 s to 0.4 s | 1.01±0.03 | 0.134±0.003 | 0.1332±0.0008 | 0.003±0.003 (0.0022±0.0006) |
| nanobeads in 60% glycerol (all) from 0.1 s to 0.4 s | 0.98±0.02 | 0.0254±0.0003 | 0.02572±0.00010 | 0.017±0.006 (0.00109±0.00007) |

**Table S4.** Results of fitting normal and anomalous diffusion models to single-particle tracking TAM[SD/$\Delta t$] data with static localization error fitted, where i is time lag. Error bars based on curve fitting, as described in Materials and methods. For $\sigma$, values without parentheses indicate fit to the anomalous model ($4\Gamma i^{\alpha}+s$) while values inside parentheses indicate fit to the linear model ($4Di+s$).

| | α | Γ ($\mu m^2/s^{\alpha}$) | D ($\mu m^2/s$) | s ($\mu m^2/s$) |
|---|---|---|---|---|
| nanobeads in peroxide (all) from 13 to 60 | 1.01±0.09 | 0.5±0.2 | 0.555±0.011 | 0±5 (0±1) |
| nanobeads in water (all) from 13 to 60 | 1.05±0.09 | 0.4±0.2 | 0.512±0.010 | 4±5 (1.2±1.0) |
| nanobeads in 40% glycerol (all) from 30 to 140 | 1.00±0.02 | 0.13±0.02 | 0.1336±0.0007 | 0.5±0.7 (0.54±0.14) |
| nanobeads in 60% glycerol (all) from 30 to 140 | 0.98±0.02 | 0.028±0.002 | 0.02529±0.00009 | 0.24±0.11 (0.38±0.02) |

**Table S5.** Results of fitting unimodal and bimodal diffusion models to single-particle tracking jump length distribution data, where i is time lag.

| | D ($\mu m^2/s$) |
|---|---|
| hemozoin in 15% peroxide (all) | i=1: 2.8±0.2<br>($D_1$) i=1: 38±6%: 0.9±0.2<br>($D_2$) i=1: 62±6%: 5.8±0.9 |
| synthetic hemozoin in 15% peroxide (all) | i=1: 0.82±0.09<br>($D_1$) i=1: 72±14%: 0.59±13<br>($D_2$) i=1: 28±14%: 4±3 |
| hemozoin in water (all) | i=1: 1.44±0.08 |
| synthetic hemozoin in water (all) | i=1: 1.06±0.06 |
| hemozoin in 40% glycerol (all) | i=1: 0.66±0.04 |
| hemozoin in 60% glycerol (all) | i=10: 0.174±0.006 |
| hemozoin in food vacuole (len<=300) | i=10: 0.37±0.02<br>($D_1$) i=10: 33±2%: 0.132±0.009<br>($D_2$) i=10: 67±2%: 0.63±0.03 |

**Table S6.** Parameters of Brownian dynamics simulations using hemozoin-like bricks or spheres. Green highlighted boxes show physical parameters of experimental expanded vacuoles.

| | # of particles | Radius of boundary (μm) | Viscosity (cP) |
|---|---|---|---|
| **Expanded FV 1** (experimental) | 35 | 3 | * |
| **Expanded FV 2** (experimental) | 23 | 2.35 | * |
| **Expanded FV 3** (experimental) | 12 | 2.3 | * |
| **Expanded FV 4** (experimental) | 16 | 1.9 | * |
| **Expanded FV 5** (experimental) | 19 | 2.4 | * |
| **Expanded FV 6** (experimental) | 11 | 1.95 | * |
| **Expanded FV 1** (brick simulation) | 35 | 3 | 60% glycerol: 10.82 Water: 1 |
| **Expanded FV 2** (brick simulation) | 23 | 2.35 | 60% glycerol: 10.82 Water: 1 |
| **Expanded FV 3** (brick simulation) | 12 | 2.3 | 60% glycerol: 10.82 Water: 1 |
| **Expanded FV 4** (brick simulation) | 16 | 1.9 | 60% glycerol: 10.82 Water: 1 |
| **Expanded FV 5** (brick simulation) | 19 | 2.4 | 60% glycerol: 10.82 Water: 1 |
| **Expanded FV 6** | 11 | 1.95 | 60% glycerol: |

| | | | |
|---|---|---|---|
| (brick simulation) | | | 10.82<br>Water: 1 |
| **Expanded FV**<br>(sphere simulation) | 40 | 2.9 | 60% glycerol: 10.82<br>40% glycerol: 3.72<br>Water: 1 |

* experimental condition, undefined viscosity

**Table S7.** Results of fitting anomalous and normal diffusion models to TAMSD data obtained via Brownian dynamics simulations of hemozoin under confinement.

| | $\alpha$ | $\Gamma$ ($\mu m^2/s^\alpha$) | D ($\mu m^2/s$) |
|---|---|---|---|
| hemozoin in water (all) from 0.1 s to 0.4 s | 0.854±0.010 | 0.704±0.011 | 0.873±0.003 |
| hemozoin in 60% glycerol (all) from 0.1 s to 0.4 s | 0.981±0.010 | 0.0947±0.0015 | 0.0976±0.0006 |

**Table S8.** Results of fitting anomalous and normal diffusion models to TAMSD data obtained via Brownian dynamics simulations of spheres under confinement.

| | $\alpha$ | $\Gamma$ ($\mu m^2/s^\alpha$) | D ($\mu m^2/s$) |
|---|---|---|---|
| Yukawa-interacting spheres in food vacuole of water viscosity from 0.1 s to 0.4 s | 0.90±0.02 | 0.88±0.03 | 1.067±0.010 |
| WCA-interacting spheres in food vacuole of water viscosity from 0.1 s to 0.4 s | 0.90±0.02 | 0.82±0.04 | 1.004±0.009 |
| LJ-interacting spheres in food vacuole of water viscosity from 0.1 s to 0.4 s | 0.89±0.02 | 0.83±0.03 | 1.019±0.009 |
| Yukawa-interacting spheres in food vacuole of 3.72 * water viscosity from 0.1 s to 0.4 s | 0.93±0.02 | 0.272±0.010 | 0.310±0.002 |
| WCA-interacting spheres in food vacuole of 3.72 * water viscosity from 0.1 s to 0.4 s | 0.93±0.02 | 0.266±0.010 | 0.302±0.003 |
| LJ-interacting spheres in food vacuole of 3.72 * water viscosity from 0.1 s to 0.4 s | 0.96±0.02 | 0.275±0.011 | 0.300±0.003 |
| Noninteracting spheres in food vacuole (R = 2.9 μm) of 10.82 * water viscosity WITH PEROXIDE | 0.94±0.03 | 0.309±0.012 | 0.333±0.002 |

| | | | |
|---|---|---|---|
| BOOST<br>from 0.1 s to 0.4 s | | | |
| Noninteracting spheres in food vacuole (R = 1.2 μm) of 10.82 * water viscosity WITH PEROXIDE BOOST<br>from 0.1 s to 0.4 s | 0.78±0.03 | 0.188±0.006 | 0.250±0.002 |
| Noninteracting spheres in food vacuole (R = 1.1 μm) of 10.82 * water viscosity WITH PEROXIDE BOOST<br>from 0.1 s to 0.4 s | 0.71±0.02 | 0.163±0.006 | 0.236±0.002 |
| Yukawa-interacting spheres in food vacuole of 10.82 * water viscosity<br>from 0.1 s to 0.4 s | 0.97±0.02 | 0.106±0.004 | 0.1110±0.0009 |
| WCA-interacting spheres in food vacuole of 10.82 * water viscosity<br>from 0.1 s to 0.4 s | 0.97±0.02 | 0.103±0.004 | 0.1090±0.0009 |
| LJ-interacting spheres in food vacuole of 10.82 * water viscosity<br>from 0.1 s to 0.4 s | 0.95±0.02 | 0.099±0.003 | 0.1085±0.0008 |

**Legends for Movies S1-S10** (all movies uploaded to figshare at the following url: https://doi.org/10.6084/m9.figshare.28835870.v2)

**Movie S1**. *P. falciparum*-infected human RBC. This video shows hemozoin tumbling from Fig. S1A. Hemozoin crystals are tumbling within the singular food vacuole (FV) compartment inside a parasite that has infected a red blood cell (RBC). Total elapsed time: 5 s. Total frame area is 8.3μm x 8.3μm.

**Movie S2**. *P. falciparum*-infected human RBC that was lethally-dosed with 0.1 μM chloroquine (see Methods). This video shows static hemozoin crystals from Fig. S1A. Total elapsed time: 5 s. Total frame area is 11.8 μm x 11. 3μm.

**Movie S3**. *P. falciparum*-infected human RBC used in TAMSI analysis. This video shows moving hemozoin in the FV that is quantified for TAMSI in Fig. S2A. Total elapsed time: 5 s. Total frame area is 10.8 μm x 9.8 μm.

**Movie S4**. Expanded FV after hypotonically treating parasites for 20 min. This video shows tumbling crystals that have enough space to spread out after FV expansion. Total elapsed time: 10 s. Total frame area is 8.0 μm x 7.2 μm.

**Movie S5**. Simulation of 35 cuboidal particles confined in a 3 μm radius spherical boundary with a viscosity of 10 cP. This video shows the Brownian dynamics of simulated hemozoin crystals in a crowded environment with the repulsive term of the Lennard-Jones potential to enforce volume-exclusion between particles. Total elapsed time: 5 s. X-, y- and z-planes are measured in m.

**Movie S6**. Simulation of 40 spheres in 2.9 μm radius spherical boundary with a viscosity of 10 cP. This video shows the Brownian dynamics of simulated spheres in a crowded environment with interaction potentials mimicking experimental potentials via the Direct Boltzmann Inversion method. Total elapsed time: 20 s. X-, y-, and z-planes are measured in μm.

**Movie S7**. Example #1 of a photosensitized parasite before light activation. Total elapsed time: 5 s. Total frame area is 10 μm x 10.8 μm.

**Movie S8**. Example #1 of a photosensitized parasite after light activation. Total elapsed time: 5 s. Total frame area is 9.9 μm x 10.9 μm.

**Movie S9**. Example #2 of a photosensitized parasite before light activation. Total elapsed time: 5 s. Total frame area is 10.3 μm x 10.1 μm.

**Movie S10**. Example #2 of a photosensitized parasite after light activation. Total elapsed time: 5 s. Total frame area is 10.4 μm x 10 μm.
